# FOXL2 interaction with different binding partners regulates the dynamics of ovarian development

**DOI:** 10.1101/2023.04.14.536403

**Authors:** Roberta Migale, Michelle Neumann, Richard Mitter, Mahmoud-Reza Rafiee, Sophie Wood, Jessica Olsen, Robin Lovell-Badge

## Abstract

The transcription factor FOXL2 is required in ovarian somatic cells for female fertility. Differential timing of *Foxl2* deletion, in embryonic versus adult mouse ovary, leads to distinctive outcomes suggesting different roles across development. Here, we comprehensively investigated FOXL2’s role through a multi-omics approach to characterise gene expression dynamics and chromatin accessibility changes, coupled with genome-wide identification of FOXL2 targets and on-chromatin interacting partners in granulosa cells across ovarian development. We found that FOXL2 regulates more targets postnatally, through interaction with factors regulating primordial follicle activation (PFA) and steroidogenesis. Deletion of one interactor, Ubiquitin specific protease 7 (USP7), induces PFA blockage, impaired ovary development and sterility. Our datasets constitute a comprehensive resource for exploration of the molecular mechanisms of ovarian development and causes of female infertility.

## Main Text

The gonads develop from bipotential genital ridges, which contain undifferentiated cell types poised to develop into ovaries in females or testes in males (*1*). The presence or absence of the Y-linked testis determination gene *Sry* determines which type of gonad it will become (*2*). Its expression is sufficient to induce testis differentiation in an XY or XX context (*3, 4*). This is also the case for its downstream target *Sox9* (*5*).

Ovarian development was considered as a default pathway activated in the absence of *Sry*, but this notion was challenged by data showing that sex determination is an active process, being governed by antagonistic signals that promote the male or female pathway and silence the alternative one (*6–8*). However, the ovarian pathway does not seem to have a clear-cut hierarchy, operating instead as a complex network of genes with considerable redundancy. *Foxl2*, a gene encoding for the forkhead box L2 (FOXL2) transcription factor is one such critical player and may have multiple roles in ovary morphogenesis, being expressed in the supporting cells of the ovary, namely granulosa cells (*9, 10*).

Previous work has highlighted the importance of FOXL2 in many aspects of female sexual development (*11*) as well as in granulosa cell-related pathologies (*12*). Heterozygous mutations in the *FOXL2* gene in humans have been linked to premature ovarian insufficiency (POI) and infertility as part of the Blepharophimosis, ptosis, and epicanthus inversus syndrome (*13–15*). Murine models have helped to pinpoint the role of FOXL2 in this syndrome. *Foxl2^−/−^*ovaries become dysgenic from 1 week postpartum, granulosa cells fail to initiate the primordial follicle activation (PFA), resulting in infertility. Mutant ovaries showed downregulation of somatic sex markers of folliculogenesis and upregulation of the pro-testis factor *Sox9* as well as others male gonadal-specific markers including *Fgf9*, *Dmrt1* and *Gata4* (*16, 17*). However, this only happens postnatally, indicating that FOXL2 is not critical for embryonic ovary specification.

In contrast, the conditional deletion of *Foxl2* from adult ovaries resulted in rapid transdifferentiation of granulosa cells into Sertoli cells, their opposite sex counterpart (*7*). This study demonstrated the plasticity of the adult ovary and the crucial role played by FOXL2 in ovarian sex maintenance. More recently, a similar role was discovered for TRIM28, which we showed to be recruited on chromatin together with FOXL2 (*6*). Collectively these data support a differential role played by FOXL2 depending on the developmental stage, as well as suggesting the presence of additional factors cooperating with FOXL2 in ensuring correct ovarian development and function.

We postulate that the multiple roles of FOXL2 during development may be dependent on the availability of different interacting partners. To address this, we performed a ChIP-SICAP analysis to identify proteins that localise with FOXL2 on chromatin and genome-wide target genes, then integrated these with RNA-Seq and ATAC-Seq data on *Foxl2* expressing cells purified from a *Foxl2^EGFP^* mouse line across ovarian development. Our data indicate a more prominent role of FOXL2 in postnatal stages. Our integrative approach uncovered a novel interactor with a role in folliculogenesis and gonadal development: Ubiquitin specific protease 7 (USP7). We show that USP7 is necessary for proper folliculogenesis, as conditional deletion in granulosa cells leads to blockage of PFA in the pre-pubertal ovary and ovarian degeneration in the adult.

### ChIP-SICAP reveals differential genomic binding of FOXL2 throughout ovarian development

To profile the dynamic changes in FOXL2 DNA binding and to identify its on-chromatin protein interactors, we performed a modified version of ChIP-SICAP (Methods & (*18, 19*)) on whole mouse ovaries at three timepoints: E14.5, one week (1W) and eight weeks (8W) postnatally (Fig.1A). ChIP-SICAP is a dual approach which allowed to simultaneously identify genome-wide FOXL2 binding sites, via ChIP-Sequencing, and proteins colocalising with FOXL2 on chromatin, via mass spectrometry. E14.5 was chosen because this embryonic stage is characterised by high FOXL2 expression in the pre-granulosa cells of the ovarian medulla (*20*); 1W coincides with the activation of primordial follicles into primary follicles (PFA), and corresponds to the timepoint at which SOX9 protein is first detected in *Foxl2^−/−^* ovaries (*10*); at 8W FOXL2 is widely expressed in the follicles where it regulates folliculogenesis (*11*) and maintains ovarian cell fate (*7*). Principal-component analysis (PCA) of ChIP-Seq peaks highlights the differences between FOXL2 binding sites across the timecourse (Fig.1B). PC1 showed a clear separation of the E14.5 samples from both 1W and 8W. PC2 further separated the 1W and 8W samples. Peak annotation showed that embryonic and postnatal FOXL2 binding sites differed in their genomic location: most identified at E14.5 (50%) were mapped to promoters, whilst only about 30% of peaks identified at 1W and 8W were located within promoters (Fig.S1A). Motif enrichment analysis of consensus peaks confirmed enrichment of FOXL2 and other forkhead factors motifs (Fig.S1B). Furthermore, we found enrichment of steroid hormone receptors binding sites amongst the top 10 predicted motifs. These included those for estrogen and androgen receptors (ESR1, *ESRRB,* AR), known to regulate hormone production and granulosa cell proliferation throughout ovarian development (*21, 22*), as well as Steroidogenic factor 1 (NR5A1), a master regulator of many aspects of gonadal development including steroidogenesis (*23*), previously identified as FOXL2 binding partner *in vitro* (*24*) and essential for gonadal development in both sexes (*25*).

**Fig.1:**
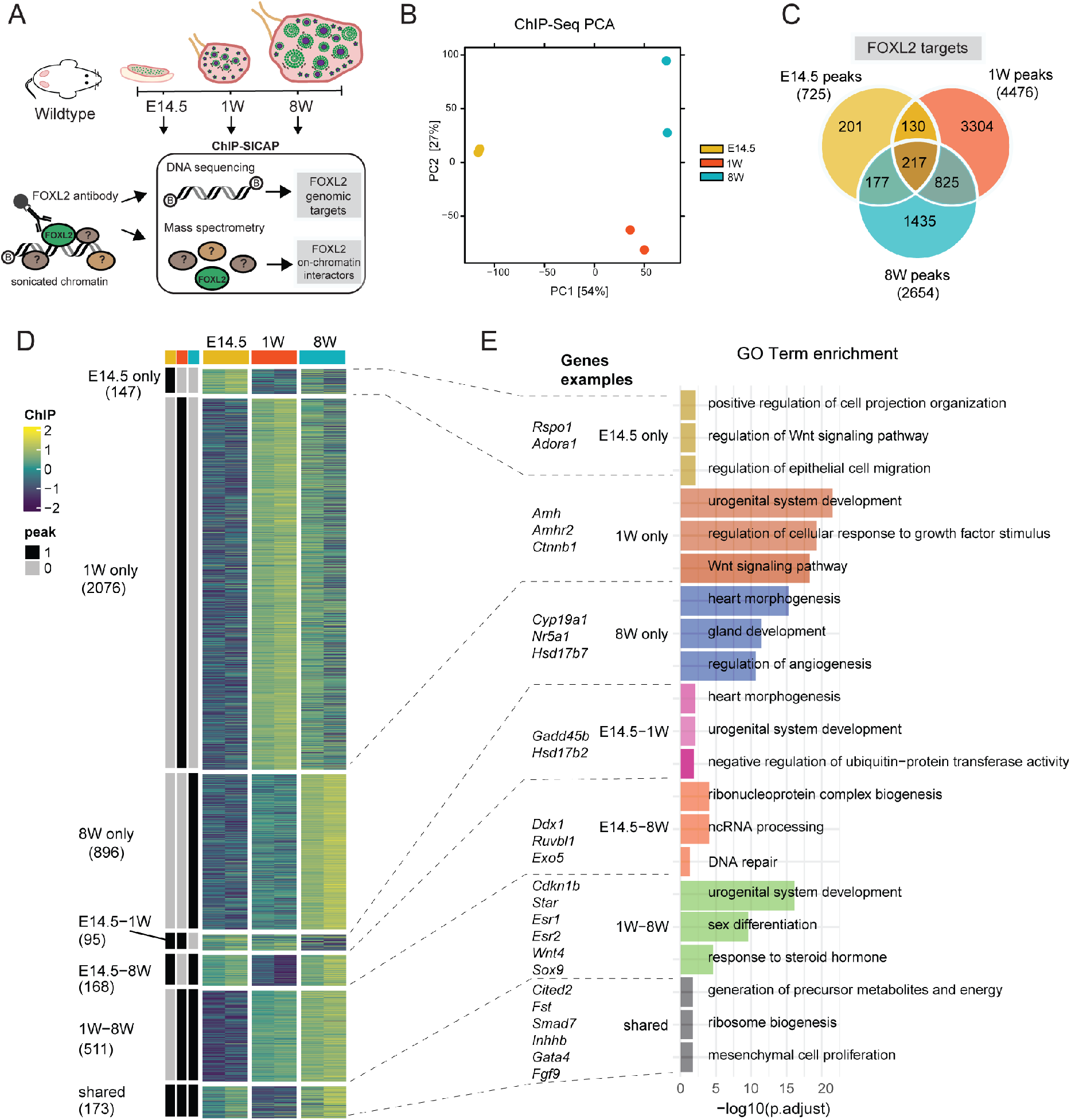
ChIP-SICAP timecourse reveals differential FOXL2 DNA binding across ovarian development. **(A)** Experimental design of FOXL2 ChIP-SICAP performed on mouse ovaries collected at E14.5, 1W and 8W, n=2. A FOXL2-specific antibody was used to capture genomics fragments bound by FOXL2 as well as proteins colocalising with FOXL2 on chromatin. (B) PCA of FOXL2 ChIP-Seq consensus peaks. (C) Venn diagram depicting the number of peaks identified at each timepoint and their overlap. (D) ChIP peak occupancy scores (gene-level). Heatmap splitting is based on ChIP peak presence/absence as indicated by black/grey bars (left). Rows are scaled by z-score. The number of genes per cluster is shown below the cluster name. (E) Bar chart showing representative GO biological processes.

In total, we detected 725 peaks at E14.5, 4476 at 1W and 2654 at 8W (Fig.1C, Data S1). Amongst all FOXL2 targets, we identified known ones bound throughout the timecourse, such as *Cited2* (*26*) (Fig.S1C), as well as targets dynamically bound including *Esr2* (Fig.S1D). Moreover, novel potential target genes were uncovered, such as *Rspo1* (Fig.S1E), a positive effector of β-catenin signaling that is required for ovarian development (*27*). Pairwise differential binding analysis allowed the identification of FOXL2-binding events that were statistically different between timepoints (Data S1).

We detected the greatest number of differential binding (2503 differential peaks) when comparing E14.5 with 8W, followed by the comparison between E14.5 and 1W (2098 differential peaks) and lastly by the comparison between 1W and 8W (634 differential peaks). By integrating our FOXL2 ChIP-Seq dataset with a microarray analysis of genes differentially expressed at 13.5, 16.5 dpc and at birth in wildtype controls versus mouse ovaries lacking *Foxl2* (*Foxl2*^-/-^, Ortiz *et al.*(*28*), Data S1), we aimed to discriminate between activation and repression activity of FOXL2 on the targets we identified. 770 genes were classified as more likely to be activated by FOXL2, because they were downregulated in the absence of *Foxl2* as well as bound by FOXL2 (Fig.S2A).

Besides *Cited2* and *Esr2*, this group included the cell-cycle repressor gene *Cdkn1b* (Fig.S2B), a known FOXL2 target that regulates granulosa cell proliferation (*29*), *Smad7* which encodes for a regulator of primordial follicle activation (PFA) together with SMAD3 (*30*) (Fig.S2C), and the follistatin gene (*Fst*) which encodes a secreted inhibitor of the TGFβ pathway, essential for ovarian development (*31*) (Fig.S2D).

A total of 376 genes were classified as more likely to be repressed by FOXL2. This group included genes encoding for signaling molecules crucial for ovary development such as *Rspo1*(*32*) (Fig.S1E), and *Wnt4*(*33*) (Fig.S2E) and exhibiting a stage-specific repression, as well as genes important for testis development such as *Gadd45g*(*34*) (Fig.S2F), and *Inhbb* (Fig.S2G)(*35*). Lastly, we evaluated FOXL2 binding in the genomic region containing *Sox9*, a gene critical for Sertoli cell and testis development(*5, 36*) (Fig.S2H). We did not detect significant binding of FOXL2 to the *Sox9* promoter at any timepoint. We then scanned the 1 Mb gene desert upstream of *Sox9*, which was previously found to contain cis-regulatory elements essential for *Sox9* expression in Sertoli cells(*37, 38*). We detected one significant peak at E14.5, two at 1W and eight at 8W (Fig.S2I). As previously shown(*6, 39*), neither TESCO enhancer nor enhancer 13 were significantly bound by FOXL2. Instead, we observed a dynamic binding of FOXL2 on the other known *Sox9* published enhancers(*37*). For example, FOXL2 was bound to Enh33 at 1W and to Enh32 at E14.5. Significant binding events of FOXL2 to the ovary-specific *Sox9* Enh8(*37*), *Sox9* Enh2 and *Sox9* Enh6 were detected at 8W. The remaining five FOXL2 binding events, four of which were detected at 8W and one at 1W, were distributed in other regions, not previously identified as Sertoli-specific *Sox9* enhancers. These may constitute granulosa-specific enhancers of *Sox9* used by FOXL2 to repress this gene in the ovary. We then clustered the ChIP-Seq peaks into sets that showed similar patterns of variation in FOXL2 occupancy across the timecourse (Fig.1D) and performed Gene Ontology (GO) analysis on each cluster (Fig.1E, Data S1) to investigate which biological processes FOXL2 regulates across ovarian development.

FOXL2 target genes known to be important in gonadal development were identified in each cluster. Those classified as shared included granulosa-enriched genes *Cited2*, *Smad7*, *Fst*, *Thbs1*, and the Sertoli-specific genes *Inhbb* and *Fgf9*. *Rspo1*, which encodes for a secreted signaling molecule that regulates WNT4 expression and is essential for female sex determination (*40*), was uniquely regulated by FOXL2 at E14.5, together with other genes including *Adora1*, *Pdgfc*, *Bcl9* and *Tert*. Those were enriched in cell migration and proliferation, cell projection organisation and Wnt signaling. Amongst the targets significantly bound by FOXL2 at 1W, we found genes involved in urogenital system development including *Amh*, *Fgfr2*, and *Ctnnb1*.

Of those genes uniquely bound by FOXL2 at 8W, we found an enrichment in those implicated in steroidogenesis pathways, including *Cyp19a1* encoding for aromatase (*41*), a known FOXL2 target (*39*), *Nr5a1*, encoding for SF1 and found previously as FOXL2 partner (*24*), and *Hsd17b7*, involved in estradiol synthesis. Whilst the cluster E14.5-1W shared several targets involved in urogenital development, the 1W-8W group featured genes encoding for members of the steroidogenesis pathway, including the known FOXL2 targets *Esr2*, for which we detected binding in the functional enhancer “Peak 3” (*39*) (Fig.S1D), the human steroidogenic acute regulatory *Star* gene (*42*) and *Cyp17a1*. Additionally, the 1W-8W included sex differentiation genes such as *Wt1* (*43*), *Esr1* (*44*), *Irx3* (*45*) and *Wnt4* (*33*), as well as Sertoli-specific genes, such as *Sox9* (*36*), *Inha* (*46*), *Fndc3a* (*47*). Lastly, the E14.5-8W cluster was enriched for DNA repair pathways, a recently discovered function of FOXL2 (*48*), and included genes such as *Ddx1*, *Ruvbl1*, and *Exo5*.

Next, we assessed the relative contribution of FOXL2 to the regulation of granulosa *versus* Sertoli-enriched genes. To obtain genes enriched in granulosa or Sertoli cells relevant for the embryonic and the postnatal stages of our timecourse, we used datasets from two studies (Fig.S3A-B, Data S2): one by Jameson et *al.* reporting microarray analysis of sorted XX (*Sry^EGFP^*) and XY (*Sox9^ECFP^*) supporting cells collected at E13.5 (*49*), and another by Lindeman *et al.* reporting RNA-Seq analysis of primary granulosa cells from mice aged 23-29 days compared to Sertoli cells isolated from *CAG-Stop^flox-tdTomato^;DhhCre* P7 XY pups (*50*), which constitute a reference for our postnatal timepoints. In this model, *Dhh-Cre* activity is specific to Sertoli cells from E12.5, therefore allowing to target specifically this cell population early on (*51*).

We calculated the overlap between FOXL2 targets at each stage of ovarian development and the granulosa-and Sertoli cell-enriched genes identified in these studies at either E13.5 (Fig.S3C), or postnatally (Fig.S3D-E). We observed that at E14.5 only 6% of FOXL2 targets corresponded to either granulosa or Sertoli-enriched genes, whilst this percentage increased to 15% at later timepoints. These comparisons, together with the greater number of targets identified at 1 and 8W compared to E14.5 in our timecourse analysis, support a greater role played by FOXL2 in controlling supporting cell-specific genes postnatally.

### Identification of FOXL2 on-chromatin protein interactors

We next used ChIP-SICAP to identify on-chromatin protein interactors (chromatome) of FOXL2 by applying mass-spectrometry on chromatin samples isolated at different stages of ovarian development (Fig.1A). Proteins co-localising with FOXL2 on the same chromatin fragments were pulled down using a specific antibody against FOXL2 followed by DNA biotinylation to separate chromatin-bound proteins.

The two-step purification strategy improves the purity of the samples by reducing the artificial interactions that are generated during the immunoprecipitation procedure and removes non-chromatin bound proteins. We identified a total of 133 proteins at E14.5, 441 at 1W, and 262 at 8W (>2-fold change FOXL2/no antibody control, n=2, adj-*pvalue*<0.1, Fig. 2A and Data S3).

**Fig.2:**
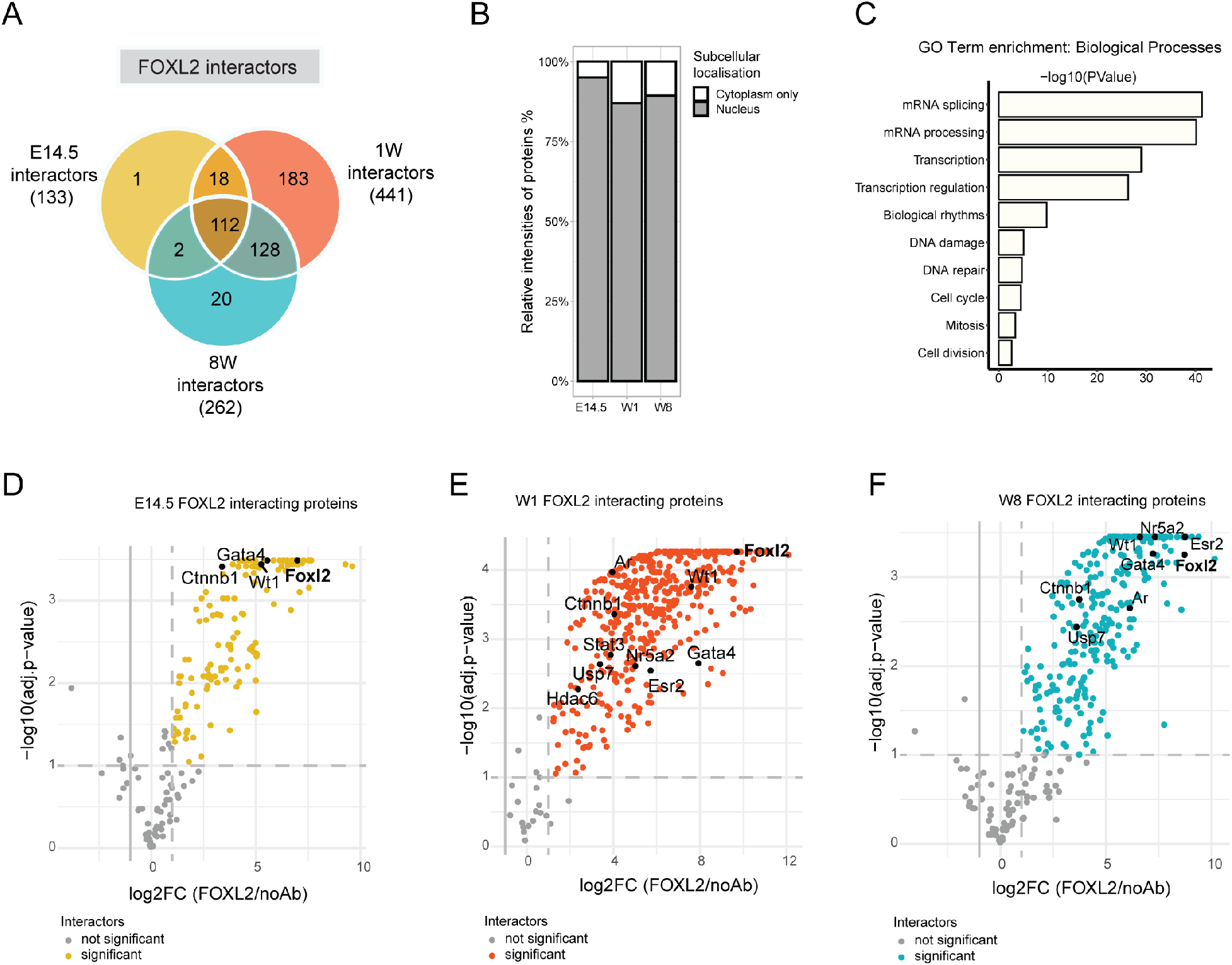
Identification of FOXL2 on-chromatin interactors by ChIP-SICAP. **(A)** Venn diagram representing the FOXL2 interactors identified by FOXL2 ChIP-SICAP. Total number of proteins is reported between round brackets. (B) Bar chart showing the relative intensities calculated using iBAQ (intensity Based Absolute Quantification) of the proteins identified and classified as either nuclear or cytoplasmatic only (proteins shuttling between nucleus and cytoplasm are classified as nuclear). (C) GO Biological processes enrichment analysis. (D-F) Volcano plot showing enrichment (mean fold enrichment >2 and adj. p value <0.1, FOXL2 immunoprecipitate *vs* no antibody control, n=2/timepoint) of proteins co-localising with FOXL2 on chromatin at E14.5 (D), 1W (E), and at 8W (F). Dotted lines separate significant proteins, also highlighted in yellow, orange and blue. Black dots: selected FOXL2 interactors.

Comparison of the relative protein intensities showed that the most highly abundant proteins were known nuclear proteins, indicating a successful enrichment for nuclear factors over potential cytoplasmic contaminants (Fig. 2B). GO enrichment analysis of FOXL2 interactors showed enrichment of proteins involved in splicing, transcription, DNA replication and repair, and cell cycle (Fig. 2C and Fig.S4). FOXL2 was amongst the top 20 mostly enriched proteins identified (based on log2FC, Fig. 2D-F, Data S3). We also identified previously known FOXL2 nuclear interactors namely STAT3, SUPT16H, ATRX, SMARCD2, XRCC6 (*52*). Amongst the FOXL2 protein partners present across the timecourse we found CTNN1B (β-catenin), a member of the WNT signaling pathway controlling the differentiation of the mouse ovary, Wilms’ tumour suppressor (WT1), a transcription factor essential for the development of the bipotential genital ridge and for the formation of the adult reproductive systems in both sexes (*53, 54*), factors important for early gonadal development and folliculogenesis such as GATA4 (*55, 56*), cell cycle regulators such as DNMT3A, heterochromatin-interacting proteins such as CBX3 and members of the splicing system including the DNA helicase DHX9, SRSF1, and DDX17. Throughout the timecourse, we also detect the E3-SUMO ligase TRIM28 which we previously showed to cooperate with FOXL2 in maintaining the ovarian cell fate (*6*).

Importantly, we identified a series of FOXL2-interacting proteins, with well-recognised roles in ovarian development and PFA in humans and mice, being differently present across the timecourse (Fig. 2D-F). We detected NR5A2 (*57*) co-localising with FOXL2 at 1W and 8W. Additionally, we found two known regulators of the primordial-to-primary follicle transition at 1W, namely the deacetylase HDAC6 (*58*) and the transcription factor STAT3 (*59*).

Androgen receptor (AR), a factor that contributes greatly to normal follicle development (*21*), was enriched at 1W and 8W but not earlier at E14.5. Importantly, we identified estrogen receptor 2 (ESR2) amongst the FOXL2 interactors at both postnatal timepoints, in agreement with the previously postulated interplay between these two factors in orchestrating key functions in ovarian physiology such as steroidogenesis (*39*) and cell fate maintenance (*7*).

We then compared our *in vivo* FOXL2 chromatome with a FOXL2 co-immunoprecipitation, proteomics dataset performed on the whole-cell granulosa-like cell line AT29C (*52*) to assess the level of overlap between our chromatin-specific dataset and one that would also include cytoplasmic interactors. We found that 112 out of the 464 proteins we identified throughout the timecourse were also found in this study. Of these shared FOXL2 interacting partners, 36 were found in our E14.5 timepoint, 109 at 1W and 63 at 8W (Fig.S4B, Data S3). These shared proteins included the known FOXL2 interactor involved in sex maintenance TRIM28 (*6*), chromatin remodelling enzymes such as ATRX, which is involved in sexual differentiation (*60*), regulators of the WNT signaling pathway such as CXXC5 and USP7, and DNA repair proteins such as XRCC6, previously identified in a yeast-two-hybrid screen and co-immunoprecipitation study (*61*). In summary, we showed that FOXL2 differentially interacts with proteins playing a role in a range of gonadal-essential functions during ovarian differentiation, from gonadogenesis, through PFA, to steroidogenesis.

### Generation of a Foxl2^EGFP^ mouse line to isolate granulosa cells

We next sought to analyse the transcriptome and chromatin landscape of granulosa cells expressing FOXL2, to ultimately integrate these datasets with the ChIP-Seq data, thus reconstructing the gene regulatory networks (GRNs) at play throughout ovarian development. We used CRISPR-Cas9 genome editing to generate a mouse line expressing an EGFP reporter knocked in the *Foxl2* locus, which allowed the isolation of cells expressing functional FOXL2 (Fig.3A, C, Fig.S5). Ovaries heterozygous for *Foxl2^EGFP^* (*Foxl2^EGFP^*^/+^) imaged by fluorescence microscopy showed a bright GFP signal specifically in the ovaries at all stages examined (Fig.3B). As expected, given the known absence of FOXL2 expression in the testis, no GFP signal was detected in the gonads of XY *Foxl2^+^*^/+^ or *Foxl2^EGFP/+^* males and *Foxl2^+/+^* XX control ovaries (Fig.3B, S5G, S6A). Colocalization of EGFP with the endogenous FOXL2 protein in *Foxl2^EGFP/+^*ovaries was confirmed by immunofluorescence (Fig.S6A). The line was maintained as heterozygotes. Fertility performance of *Foxl2^EGFP/+^*females was monitored over 12 weeks and found to be normal compared to control females (Fig.S5H).

**Fig.3.**
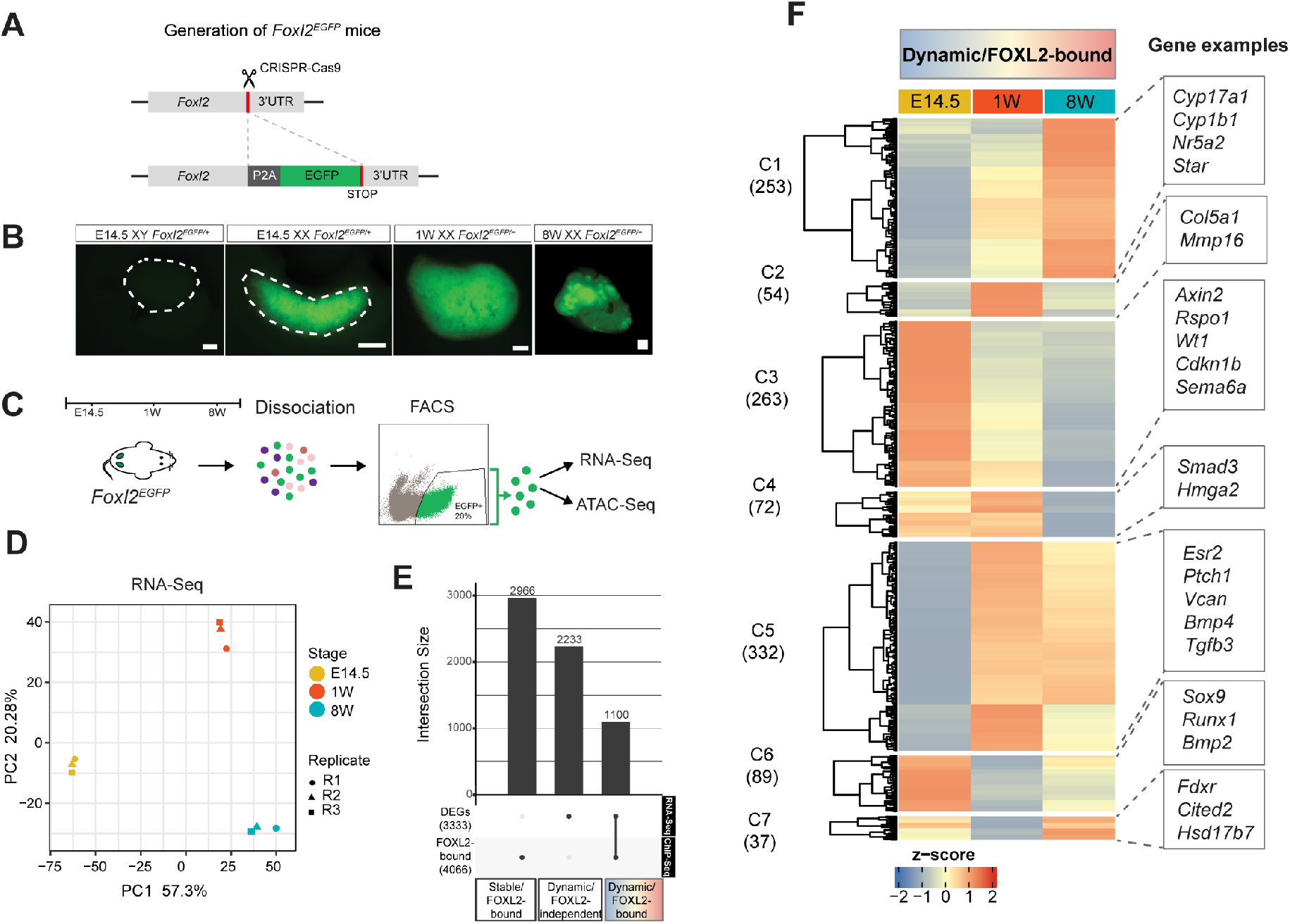
An integrative approach combining RNA-Seq on *Foxl2^EGFP^* granulosa cells and ChIP-Seq refines the gene regulatory networks controlled by FOXL2. **(A)** CRISPR/Cas9 mediated EGFP insertion at the *Foxl2* locus. (B) Representative images of EGFP fluorescence in freshly collected gonads. Scale bars: 200 µm. Dotted lines outline the embryonic testis (left) and ovary (right). (C) Overview of sample collection and downstream analyses. (D) PCA plot of RNA-Seq analysis of cells isolated from E14.5, 1W and 8W gonads (n=3). (E) Number of differentially expressed genes (DEG) identified by RNA-Seq, FOXL2 gene targets identified by ChIP-SICAP and the overlap between the two datasets. (F) Hierarchical clustering of the 1100 genes bound by FOXL2 and differentially expressed across the timecourse. Gene examples are shown on the right.

We dissociated ovaries from *Foxl2^EGFP/+^* mice at E14.5, 1W and 8W (Fig.3C) and isolated GFP positive cells by fluorescence-activated cell sorting (FACS). A total of 50k and 100k of *Foxl2^EGFP^*positive cells were processed for bulk RNA-Seq and ATAC-Seq respectively, using three biological replicates for each timepoint.

PCA of RNA-Seq data (Fig.3D) based on the variance stabilised abundance estimates showed good separation between the timepoints indicating major transcriptional changes characterising granulosa *Foxl2^EGFP^* positive cells throughout time. Most of the variance was evident along the first principal component, which separated E14.5 samples from both W1 and W8. Poisson similarity metric indicated more similarity between the transcriptome of postnatal samples (1W and 8W) compared to E14.5 (Fig.S7A). To examine the global transcriptome dynamics, we selected genes significantly changing across the timecourse in at least one pairwise comparison (Data S4). Differential gene expression analysis comparing 8W and E14.5 revealed the greatest changes with 1436 genes upregulated and 1134 downregulated; comparing 1W with E14.5 revealed upregulation of 1307 genes and downregulation of 861 whilst the least number of changes were detected between 8W and 1W, with 489 genes upregulated and 556 genes downregulated (Fig.S7B). Because FOXL2 is also expressed in theca cells (*62*), we first assessed the expression values of known theca and granulosa cell markers (*63*). We found an enrichment of granulosa over theca markers, confirming the suitability of this dataset for the exploration of granulosa-specific GRNs (Fig.S7F). We then used k-means clustering to analyse the dynamics of gene expression changes across time (Fig.S7C-D) and identified seven main patterns. Clusters 7 and 6 consisted of genes highly expressed at E14.5 including markers of pre-granulosa cells such as *Rspo1*, *Lgr5* and *Wnt4*, important for the initial cell proliferation in XX embryonic gonads (*27*). GO term biological processes analysis (GO BP), (Fig.S7E), of these clusters revealed enrichment of biological processes such as Wnt signaling and urogenital system development pathways. Cluster 5 included genes moderately expressed at E14.5 and upregulated specifically at 1W and included genes involved in kidney and urogenital system development, genetic imprinting, and extracellular matrix organisation. Cluster 1 consisted of genes uniquely up regulated at 1W and participating in extracellular matrix organisation, cell cycle phase transition, and DNA replication. Cluster 4 grouped genes activated as the ovary develops from 1W and remain upregulated in adulthood at 8W. GO analysis showed enrichment in nuclear division, cell cycle, lipid transport, and synapses organisation. Lastly, Cluster 3 included genes specifically upregulated at 8W and enriched in steroid metabolic processes, and angiogenesis. These data show the timely activation of genes important for embryonic and postnatal ovarian development and function in our system.

### An integrative approach to reconstruct gene regulatory networks controlled by FOXL2

Having identified the gene expression changes characterising *Foxl2^EGFP/+^* granulosa cells, we then sought to reconstruct the GRN underpinned by FOXL2 (Fig.3E and Data S5).

We took the total number of FOXL2-bound genes identified by ChIP-Seq at any given timepoint (Fig.1) and intersected it with the set of DEGs identified by RNA-Seq. Genes that changed in their expression value at any point between E14.5, 1W and 8W (3333 genes) and that were assigned to a FOXL2 ChIP peak in at least one timepoint (4066 genes) were considered as “dynamic/FOXL2-bound” genes. The rest of the genes bound by FOXL2, but not changing their expression values were considered as “stable” (2966 in total) and divided between “stably repressed” and “stably activated”, based on a variance stabilised (VST) cut-off of 8.3 (Fig.S8A).

This threshold was chosen based on the expression values of known granulosa-enriched genes (*Foxl2*, *Esr2*, *Rspo1*), versus genes residing on the Y chromosome therefore not expected to be expressed at all in XX ovaries (*Sry*, *Usp9y* and *Uty*), as well as on the expression values of Sertoli-enriched genes (*Dmrt1*, *Sox9* and *Fgf9*) (Fig.S9A-C). Using this approach, we identified a total of 777 genes bound by FOXL2, but which are not expressed at any point during the timecourse and are therefore assumed to be repressed by FOXL2. These were enriched in cell-cell adhesion, immune, and protein secretion pathways (Fig.S8B). These also included Sertoli-enriched genes such as *Spz1* (*64*), *Cdh8*, *Rims1*, *Fgf9* (*65*), *Hsd17b3*, *Hopx* and *Tuba3* (*50*). In contrast, a significant higher number of genes were bound by FOXL2 and were expressed throughout the timecourse but did not significantly change in levels across this (2189 genes). These were enriched in mRNA processing, ubiquitin-dependent catabolic processes, cell cycle and histone modification pathways. In contrast, the “dynamic/FOXL2-bound” group contained a total of 1100 genes. Unsupervised hierarchical clustering identified 7 main clusters (Fig.3F). GO enrichment analysis (Fig.S8C) of these clusters showed that FOXL2 regulates a plethora of dynamically expressed genes enriched for functions critical for different stages of ovarian development. For example, we found enrichment in steroid metabolic and angiogenic processes, typical of folliculogenesis, in cluster 1 (C1), which included genes upregulated specifically at 8W. Of these, 53% were also a direct target of FOXL2 specifically at 8W (Data S5). On the contrary, if considering C3, C4 and C6, clusters included genes upregulated at E14.5 compared to 8W, only 12% of the genes were significant targets of FOXL2 at E14.5. This confirmed the preponderant role of FOXL2 in regulating significantly changing genes at postnatal stages rather than during embryonic development. The high enrichment of Wnt signaling pathway genes found in C3, which contained a total of 263 genes upregulated at E14.5 only 36 of which were bound by FOXL2 at this timepoint (including several Wnt signaling pathway effectors such as *Rspo1*, *Zfp703* and *Bcl9*), was consistent with the established role of this pathway in ovarian development (*33*).

### *Foxl2^EGFP^* positive cells have a dynamic chromatin landscape across ovarian development

To explore the dynamics of chromatin remodelling in granulosa cell development, we performed ATAC-Seq on *Foxl2^EGFP^* positive cells (Fig.4A, Data S6).

**Fig.4:**
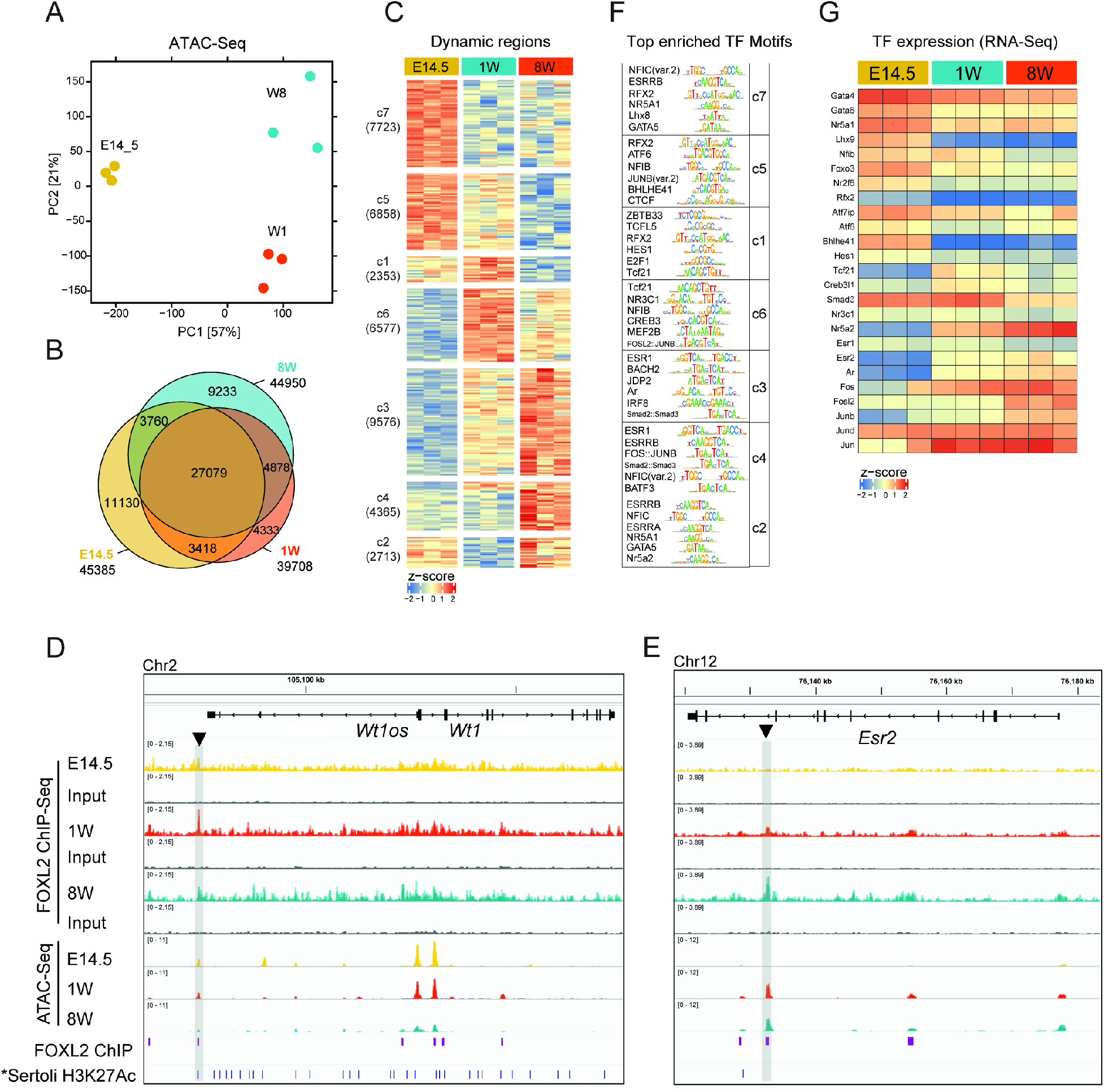
Granulosa cells have a dynamic chromatin landscape throughout development. **(A)** PCA of ATAC-Seq performed on *Foxl2 ^EGFP/+^* sorted ovarian cells at E14.5, 1W and 8W. (B) Venn diagram reporting the number of significant peaks. (C) Hierarchical clustering of peaks. (D-E) IGV snapshots of representative regions. *Sertoli H3K27Ac ChIP-Seq was re-analysed to match mm10 coordinates from Maatouk e*t al* (*67*). (F) MonaLisa analysis of TF motifs enrichment analysis within the open chromatin regions. Top 6 motifs for each cluster are reported (Table S6). (G) Heatmap displaying gene expression changes of representative TF, as assessed by RNA-Seq (Table S4).

In total, we identified 45385 accessible regions at E14.5, 39708 at 1W, and 44950 at 8W (Fig.4B, S10A). Of these, 27079 were accessible throughout the timecourse, while 11130 were unique to E14.5, 4333 unique to 1W, and 9233 unique to 8W. The genomic annotation of open chromatin regions was similar across timepoints, with 32-35% mapping to promoters, 34-38% to the gene body, 3% downstream of a gene, and 25-28 % mapping to distal intergenic regions (Fig.S10B). K-means clustering of consensus peaks revealed 7 patterns representing similar dynamic opening and closing of chromatin regions (Fig.S10C). To assess the potential contribution of FOXL2 in binding to these regions, we specifically looked for enrichment values for the FOXL2 canonical motif (JASPAR id: MA1607.1) in each ATAC-Seq cluster (Fig.S10D). This analysis revealed a greater enrichment in clusters 6, 3, 4, 2 and the least in cluster 7 (fold enrichment 1.4-1.2, adjPvalue>4). Amongst these, clusters 6 and 3, which included peaks accessible preferentially at 1W and 8W and closed at E14.5, were the two most significantly enriched with FOXL2 motifs. In contrast, amongst regions accessible at E14.5 (c1, c5 and c7) only c7 contained significant enrichment of FOXL2 binding sites, albeit with an enrichment score much lower than c6 and c3.

We performed differential accessibility analysis using DESeq2(*66*) (Data S6) and identified a total of 40165 differentially accessible regions (DARs) throughout the timecourse, 32611 of which were changing between E14.5 and 8W, 24026 were changing between E14.5 and 1W, and 16054 were changing between 1W and 8W. These chromatin landscape changes were consistent with the transcriptomic data, which showed that most of the differences were detected comparing E14.5 and 8W samples. DARs included novel potential cis-regulatory regions, for example one within the *Rspo1* gene, conserved, and differentially bound by FOXL2 according to our ChIP-SICAP analysis, and showing a similar change in its accessibility, as indicated by ATAC-Seq (Fig.S10E). We also detected FOXL2 binding by ChIP-Seq at a known *Wt1* enhancer located 50 kb upstream of its promoter (*67*) (Fig.4D). This binding correlated with closing of this region at 8W. In contrast, *Esr2* was bound by FOXL2 at its known functional enhancer (*39*) only at postnatal stages (Fig.4E). This enhancer was not marked by H3K27Ac (active enhancer mark) in Sertoli cells, therefore can be considered as a granulosa-specific cis-regulatory region of *Esr2*.

K-means clustering identified seven clusters representative of the main chromatin remodelling patterns (Fig.4C). Amongst the DARs, we detected regions located nearby granulosa cell markers such as *Fst* (*68*) and *Runx1* (*8*) which were highly conserved (Fig.S10E). Additionally, we identified open elements nearby markers of Sertoli cells, including structural markers such as *Cldn11* and *Itga6*, hormonal markers such as *Amh* and *Hsd17b1*, and genes encoding for transcription factors such as *Nr5a1* and *Fgf9*. These were also bound by FOXL2 as assessed by ChIP-SICAP.

To investigate what transcription factors may bind to these dynamic regions that could potentially drive the observed gene expression changes or that may direct changes in chromatin organisation, we performed a transcription factor (TF) motif enrichment analysis of the DAR dynamics using monaLisa (*69*) (Fig.4F, Data S5). Regions that were more accessible at E14.5, but lost accessibility at later stages (namely c7 and c5) were enriched with nuclear factor family motifs and GATA motifs. NR5A1 motifs were highly represented, as well as motifs from the RFX family. Cluster 5 was also enriched in motifs typical of the transcriptional repressor CTCF. Regions accessible at 1W (c1 and c6) were enriched in motifs for factors involved in Wnt signaling (TCFL5, TCF21) and Notch signaling (HES1) and for ZBTB33 motifs, a transcriptional repressor also known as Kaiso, which negatively regulates the Wnt signaling pathway (*70*). Amongst these, TCF21, ATF1 were also found to localise with FOXL2 on chromatin postnatally, as found by FOXL2 ChIP-SICAP (Data S3 & Data S5). Clusters containing regions opened at 8W were enriched with motifs for members of the steroid hormone receptor family, including ESRRB (containing a half-site for the binding of ESR2) and AR, factors known to play critical roles in folliculogenesis and ovulation (*21, 71*), which were also found in our proteomics analysis (Fig.2). ESRRB motifs were also enriched at E14.5 although no expression of *Esr2* mRNA was found at this timepoint by RNA-Seq (Fig.4G). Lastly, in c2 we found enrichment for the TF NR5A2, known to regulate PFA (*57*), which was also detected with FOXL2 ChIP-SICAP at 1W and 8W timepoints.

RNA-Seq analysis revealed that some of these factors also show an expression pattern matching the binding motif predictions. For example, GATA4 and LHX9 expression was increased at E14.5, whilst factors predicted to bind at postnatal stages including NR5A2, AR and ESR2, were upregulated from 1 week onwards.

### Deletion of Usp7 in granulosa cells blocks primordial follicle activation and impairs ovarian development

To select for novel players controlling PFA, we focussed on proteins that were bound with FOXL2 on chromatin at 1W and 8W, when PFA occurs, but not at E14, and that were associated with any of the following terms in OMIM database: “premature ovarian failure”, “hypogonadisms” or “infertility”, to enrich for factors more likely to be relevant to human conditions. We found one protein named USP7 (Figure 2E-F) satisfying these criteria, where individuals carrying gene mutations exhibited developmental delay, intellectual disability, and hypogonadism in both sexes (*72, 73*). USP7 is recognised to play a role in chromatin regulation directly through the depletion of ubiquitin marks involved in DNA methylation (*74*), and indirectly by controlling the stability of DNA methyltransferases (*75, 76*). USP7 is also a regulator of Wnt signaling, a pathway with a pivotal role in pre-granulosa cell activation during PFA (*77*).

To test the function of USP7 in granulosa cells, we conditionally deleted *Usp7* (*78*) using *Sf1:Cre^tg/+^*transgenic mice (*79*). *Usp7^fl/+^;Sf1:Cre^tg/+^*mice of both sexes did not display any obvious morphological or reproductive phenotypes. We then analysed the phenotype of *Usp7^fl/fl^*mice lacking the Cre driver transgene, defined herein as control, and compared it to those of *Usp7^fl/fl^;Sf1:Cre^tg/+^* mutants.

In homozygous mutants, no evident morphological abnormalities were observed when comparing ovaries to wildtype mice at P0 (Fig.5A, a-b). However, immunofluorescence analysis showed that the clear boundary between the cortex and medullary region of the ovary, present in wildtype, was lost in mutant ovaries (Fig.5A, c-d). FOXL2/DDX4 double staining revealed that in wildtype ovaries, germ cells were densely packed in the cortical region, whilst individual oocytes surrounded by FOXL2-positive granulosa cells started to organise themselves into what would later develop as individual follicles in the medulla.

**Fig.5:**
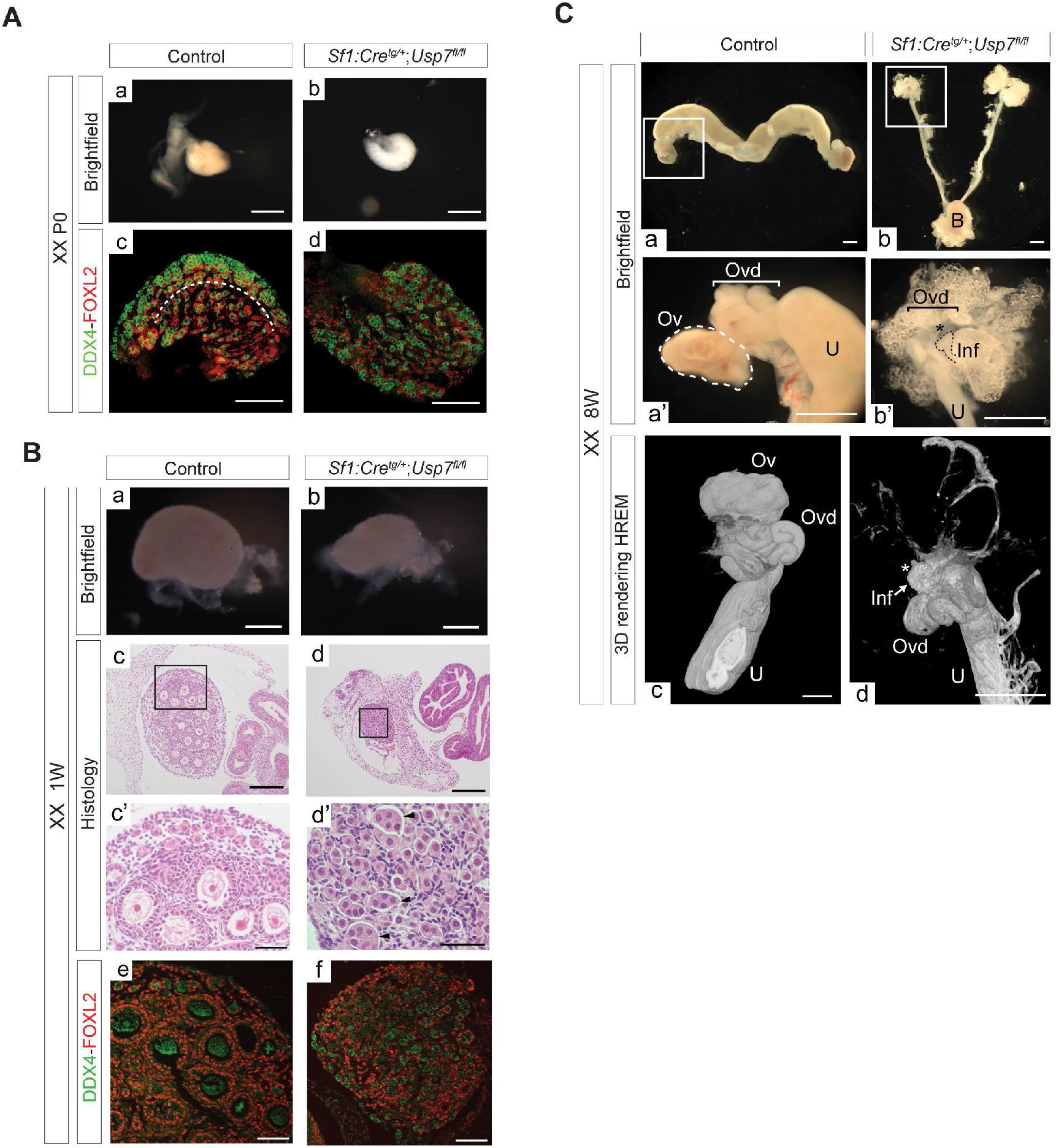
Deletion of *Usp7* in granulosa cells blocks primordial follicle activation and impairs ovarian development. Ovaries of *Usp7^fl/fl^* mice (control) and *Usp7^fl/fl^;Sf1:Cre^tg/+^* mice (mutants). Representative images of n=3 experiments. (**A**) Brightfield images showing morphology of control (a) and mutant (b) P0 ovaries, Scale bar=500µM in all, unless otherwise specified. (A, c-d) Immunofluorescence analysis of DDX4 (green) and FOXL2 (red). (B, a-b) Brightfield images of 1W ovaries. (B, c-d) H&E staining, scale=200 µM. (B, c’-d’), higher magnification showing persistence of nest-like structures in mutants (arrowheads in d’, scale=50 µM). (B, e-f) Immunofluorescence staining for DDX4 and FOXL2 in 1W ovaries. (C) Reproductive tracts and ovaries from 8W control and mutant mice. (a’-b’) Higher magnification, scale=1mm. (c-d) HREM 3D rendering of reproductive tract from a 8W control ovary and a XX mutant. Asterisk indicates the area where an ovary should be. Arrow indicates infundibulum tip and lack of ovarian structure. Scale=0.7mm. Ovd=oviduct, Ov=ovary, U=uterus, B=bladder, Inf=infundibulum.

In contrast, mutant ovaries did not display such demarcation, granulosa cells appeared disorganised in their spatial localisation, and most oocytes remained in germ cell nests.

At 1 week postnatally we observed that mutant ovaries were visibly smaller compared to controls (Fig.5B, a-b). Control ovaries contained a cortex of primordial follicles, where dormant oocytes are surrounded by one layer of flattened granulosa cells, and a medullary region, containing follicles which have been activated and progressed towards the primary follicle stage (Fig.5B, c-d). In these, granulosa cells were well organised, with some having completed the transition from flattened to cuboidal, and strongly expressing FOXL2 (Fig.5B, c’, e). Strikingly, in mutant ovaries, only primordial follicles were found (Fig.5B, d’, f). Several nest-like structures were identified encapsulating clusters of germ cells, which are not expected at this age. Granulosa cells in *Usp7^fl/fl^;Sf1:Cre^tg/+^*mice were flattened, expressed FOXL2 and appeared to be arrested at the primordial follicular stage. These data indicate that expression of USP7 in the somatic cell lineage is necessary to activate primordial follicles towards their developmental pathway, which will ultimately result in either atresia or ovulation. In adult ovaries (Fig.5C, a-b), no ovarian structures could not be found in 8 weeks old *Usp7^fl/fl^;Sf1:Cre^tg/+^* mice, and the oviduct terminated with the infundibulum tip (Fig.5C, b’ and d, Movie S1), indicating complete degeneration of the ovary in the absence of *Usp7*.

The uterus of 8 weeks old mutant mice was markedly hypoplastic, consistent with hypoestrogenism expected to occur consequently to the loss of ovarian function. Because *Usp7* was reportedly found mutated in XY patients exhibiting hypogonadism (*72, 73*), we also investigated the phenotype of mutant testes. *Usp7^fl/fl^;Sf1:Cre^tg/+^* testes were smaller than controls from P0 (Fig.S11A, a-b). Immunofluorescence staining revealed an increased number of DD4X4-positive germ cells in *Usp7^fl/fl^;Sf1:Cre^tg/+^* mice compared to control littermates (Fig.S11A, c-d). The size difference between control and mutant gonads was accentuated at 1 week postnatally (Fig.S11B, a-d), when testes started to become markedly hypoplastic. Immunofluorescence for SOX9 revealed abnormal distribution of Sertoli cells and disorganised tubules (Fig.S11B, e-j). The mutants often presented with a fluid-filled cavity lined by a SOX9-negative epithelium. The hypoplasia of mutant testes was more pronounced in adult gonads, when the testes of mutant mice exhibited a 98% decrease in their weight (Fig.S11C-left and D, a-b). Homozygous mutants were completely sterile (Fig.S11C, right). Mature elongated spermatids were absent in mutants (Fig.S11D, d-d’), whilst these were found in controls (Fig.S11D, c-c’). Tubules were dysgenic, showing a reduction in the number of Sertoli cells (SOX9, red in Fig.S11D, e-f), and often exhibiting depletion of germ cells from defective tubules (Fig.S11D, g-j, asterisk).

These results suggest that USP7 is necessary for the correct specification and function of somatic cell lineages in both the ovary and the testis, and for fertility in both sexes. FOXL2 is likely to be a critical interacting factor for USP7 in the ovary, but other partners may be present in the testis where *Foxl2* is not expressed and interact with USP7 in this context.

## Discussion

In this study, we describe the set of FOXL2 genomic targets, on-chromatin protein interactors, gene expression changes and chromatin landscape dynamics characterising granulosa cells at three critical stages of mouse ovarian development. We found that FOXL2 binds to a greater number of genes at postnatal compared to embryonic stages, supporting a more critical role for this factor during postnatal life compared to the embryo, at least in mice. GO analysis of FOXL2 targets at E14.5 showed enrichment of Wnt signaling, in keeping with the synergistic role played by these two pathways in reinforcing ovarian cell fate commitment (*80*). The Wnt pathway has been shown as essential for pre-granulosa cell activation during PFA (*77*). This pathway was also highly featured amongst FOXL2 target genes identified at 1W, supporting the idea that a possible interplay between Wnt and FOXL2 around the time of PFA may happen through regulation of similar targets. Genes bound at 8W, once mice have reached sexual maturity, are enriched for steroidogenesis and DNA repair pathways. Notably, we found FOXL2 to directly bind to a known ESR2 enhancer (*39*), specifically at 8W. Indeed, in addition to binding sites for forkhead factors, FOXL2 peaks were highly enriched in steroid receptors motifs including ESR1 and AR. This supports previous *in vitro* studies which indicated a role for FOXL2 on estradiol signaling (*39*). One of the main roles of FOXL2 in the ovary is direct repression of *Sox9* (*7*). Our data show that this repression may be time-specific, as we detected a greater number of FOXL2 peaks in the ‘gene desert’ regulatory region 5’ to *Sox9* locus at 8W, compared to earlier timepoints. However, this does not exclude that the presence of a single peak at earlier timepoints may suffice and act as repressor site for *Sox9*. In agreement with other studies, we did not find binding of FOXL2 to TESCO (*6, 11, 26*), nor to the distal *Sox9* Enh13 (*37, 81*), but we instead detected peaks in other regions, mostly not overlapping with Sertoli-specific *Sox9* enhancers (*37*). In addition, FOXL2 appeared to bind to several Sertoli-enriched genes, with structural roles (*Cldn11*, *Itga6*) or involved in hormone signaling (*Hsd17b1*, *Amh, Inhbb*). These data suggest that FOXL2 may repress testis development postnatally through alternative *Sox9* enhancers, many of which are granulosa-specific, but also by directly repressing genes important for Sertoli cell proliferation and function. The latter may well be the most important function of FOXL2 beyond repressing *Sox9*. This was suggested to be the case in the conditional TRIM28 null mutant XX gonads, where expression of Sertoli structural genes precedes *Sox9* upregulation in an intermediate cell type along the female-to-male transdifferentiation trajectory (*6*).

We then queried the nature of the binding partners that were co-localising on the same chromatin regions with FOXL2. Our ChIP-SICAP analysis represents the first *in vivo* analysis of chromatin binding partners of FOXL2 and revealed that this factor interacts with proteins crucial for each developmental milestone of ovarian development. For example, we found that FOXL2 colocalises with the transcription coactivator of Wnt signaling CTNNB1 (also known as β-catenin) throughout the timecourse. β-catenin activation was shown to induce male-to-female sex-reversal in a normal XY gonad (*82*), together with increased expression of FOXL2 and SOX9 downregulation. The co-localisation of β-catenin and FOXL2 may represent a mechanism to reinforce ovarian cell fate, suggesting that the functional interplay between the two pathways may rely on their joint recruitment to DNA. This was also supported by the identification of several other members of the Wnt pathway including USP7. USP7 is a disease-associated deubiquitinase (DUB) (*83*), playing a role in chromatin remodelling through its histone deubiquitination activity (*74, 75*), which can affect gene transcription activation and silencing (*84*). We found that USP7 is necessary to activate primordial follicles in mouse ovaries. The fact that *Usp7* mutant ovaries still express FOXL2 suggest that USP7 may regulate PFA in a redundant way with FOXL2, as previously shown for other members of the Wnt signaling pathway (*80*). Or alternatively, that USP7 may regulate the function of other players involved more directly in PFA, such as FOXL2 itself. It has recently been shown that USP7 can stabilise transcription factor activity such as that of AR, by reversing polyubiquitination, thus allowing binding to DNA (*85*). We speculate that USP7 activity might affect FOXL2 recruitment to DNA, stabilising this interaction and thus allowing the execution of the primordial-to-primary transition in wildtype prepubertal ovaries. We have recently shown that the epigenetic regulator TRIM28 interacts with FOXL2 to maintain the ovarian cell fate (*6*). TRIM28 also interacts with SOX9 in fetal testes (*86*). Our findings on USP7 add to the growing evidence supporting the role of epigenetic factors, often with bivalent roles in the two sexes, in the specification of somatic cell lineages of the gonads. We identified four other interactors known to have a role in PFA that had been presumed to be FOXL2-independent, namely NR5A2 (*57*), HDAC1 (*87*), HDAC6 (*58*) and CSNK1A1 (*88*). This indicates that FOXL2 may create a chromatin hub where factors regulating similar processes can interact. Our study constitutes a precious resource for the discovery of novel regulators of PFA, a fundamental process that must be tightly regulated to ensure long lasting reproductive fitness. Recently, *in vitro* activation, a method for controlling PFA, has provided a possibility for conceiving to POI patients (*89*). Identification of novel regulators of PFA may therefore also be invaluable to expand the toolset in reproductive technologies.

FOXL2 co-localised with WT1 throughout ovarian development. WT1 is a transcriptional regulator essential for early development of the gonads (*90*) existing in two main splicing isoforms, +KTS and -KTS. Only -KTS can act as a transcriptional regulator because the addition of the +KTS portion abrogates WT1 ability to bind to DNA, confining it to nuclear speckles(*91, 92*). Given this knowledge and the fact that our FOXL2 ChIP-SICAP experiments specifically target proteins bound to chromatin, which is typically absent from nuclear speckles (*93*), our data support an interplay on the DNA of FOXL2 and WT1 -KTS, which may aid the suggested role of the latter in early ovarian development (*94, 95*). We also identified interaction of FOXL2 with GATA4 (*56, 96*). It was previously shown that deficiency in GATA4-FOG2 interaction impaired ovarian development through the alteration of the female gene expression program, including having a direct negative effect on *Foxl2* expression (*97*). Our data indicate that interaction with FOXL2 may be at the base of GATA4’s role in ovarian development. At 1 and 8W, we found FOXL2 interacting with ESR2, confirming that the previously postulated interaction between FOXL2 and estrogen signaling is mediated through the physical interaction of ESR2 and FOXL2 at proximal genomic locations (*39, 98*). In addition, we detected AR at 1W and 8W as interactors, which supports the motif analysis showing enrichment of AR motifs in FOXL2 binding sites. AR has a dual role, regulating granulosa cell proliferation in the ovary (*21*), while it is critical to ensure spermatogenesis is supported by Sertoli cells in the testis (*99*). This co-localisation may hint to a role for AR in the transdifferentiation phenotype observed in the conditional *Foxl2* mutants (*7*). We generated a *Foxl2^EGF^*^P^ mouse line to isolate FOXL2-positive cells. This was used to produce age-matched transcriptome and chromatin landscape datasets to integrate with our ChIP-SICAP data on the genomic binding sites. This integration allowed us to identify a high confidence set of genes likely to be repressed, activated, or dynamically regulated by FOXL2 across ovarian development. Our analysis showed that the majority of FOXL2 targets were either stably expressed throughout the timecourse, or dynamic, while a lower number of genes were bound by FOXL2 and stably repressed. Amongst the stably expressed genes, we found genes with a role in mRNA processing, apoptosis and cell cycle, function which have all been linked with FOXL2 in functional studies (*100*). Analysis of those that are dynamically expressed and controlled by FOXL2 provided an overview of the range of functions critical for the ovary that this factor may regulate, from steroidogenesis to extracellular matrix organisation. Genes involved in axon guidance were also highlighted, which is of interest as the ovary is innervated, whereas the testis is not (*101*). A smaller number of genes were stably repressed, including genes encoding for Sertoli structural proteins. We identified several potential enhancers of genes important for granulosa cell differentiation, in genes such as *Rspo1* and *Runx1*. Particularly in the latter, several FOXL2 binding sites coincided with open chromatin regions, which agrees with the FOXL2-RUNX1 interplay in regulating ovarian cell fate maintenance (*8*). Finally, the integration of ATAC-Seq with FOXL2 ChIP-SICAP and comparison with ATAC-Seq datasets from sorted Sertoli cells (*50, 102*) also allowed us to observe that many genes important for Sertoli cells (e.g. *Amh*, *Cldn11*, *Itga6*, *Fgf9*, *Hsd17b1*) contained regions open in both Sertoli and granulosa cells and that these were bound by FOXL2 in granulosa cells. This suggests that the role of FOXL2 in repressing the male pathway may go beyond only repressing *Sox9* and may unfold additionally through the direct repression of genes encoding proteins critical for the structure and function of Sertoli cells. It does not exclude, however, that it is the activation of *Sox9* and *Dmrt1*, the primary event leading to the gonadal sex reversal. The bivalent state of these regulatory regions also reveals that the system is permanently ready to undergo transdifferentiation from granulosa to Sertoli cells or *vice versa*.

In summary, our study represents the first *in vivo* description of the chromatin interactome of the ovarian-specific transcription factor FOXL2. We showed that this factor, essential for female fertility, differentially interacts with proteins playing a role in a range of gonadal-essential functions, from gonadogenesis, through PFA, to steroidogenesis and that this action is stage-specific. We identified a novel regulator of primordial-to-primary follicle transition, USP7, and showed that its granulosa cell-specific deletion leads to the inability of primordial follicles to leave the arrested pool stage and progress throughout folliculogenesis.

This study provides a rich resource of novel protein interactors colocalising with FOXL2 on nearby genomic locations that could be used to generate new hypotheses on the molecular mechanisms of ovarian development, POI, granulosa cell tumours, and other causes of female infertility.

## Acknowledgments

We thank members of the Lovell-Badge lab for advice. We particularly thank Dr Silvana Guioli and Dr Karine Rizzoti for their helpful feedback on this manuscript. We are particularly grateful to Dr Nitzan Gonen (Bar-Ilan University), to Dr Francis Poulat (University of Montpellier), and to Dr Marie-Christine Chaboissier (Institut de Biologie Valrose) for scientific discussions and support. We thank the Francis Crick institute platforms (GeMS, ASF, BABS, FLOW, Proteomics, EHP, CALM) for their excellent support and for sharing their expertise. We thank Jacek Mor and all staff from the Biological Research facility (Francis Crick Institute) for taking excellent care of the animals. We thank Fabrice Prin for help with the HREM experiments. We thank Dr Dagmar Wilhelm for the generous gift of the anti-FOXL2 antibody and for valuable feedback.

## Author contributions

Conceptualization: RM

Visualization: RM, MN

Investigation: RM, MN

Bioinformatic analyses: RM, RM, MRR

Methodology: RM, MN, MRR, SW, JO

Writing-original draft: RM

Writing-review&editing: RM and all authors

Funding acquisition: RLB

Supervision: RM, RLB

## Supplementary Materials

Materials and Methods

Figs. S1 to S11

Tables S1

Movies S1

Data S1 to S6

## Supplementary Materials and methods

### Animal procedures

All animal regulated procedures carried out were approved under the UK Animals (Scientific Procedures) Act 1986 and under the project licences 80/2405 and PP8826065 and Home Office guidelines and regulations The *Foxl2^EGFP^*allele was maintained on a C57BL/6 background. *Foxl2^EGFP/+^* embryos were produced by crossing heterozygotes with C57BL/6 wildtype animals and collected at E14.5 after timed matings. Noon of the day of plug was considered as E0.5. Litters of 1-week old postnatal mice were sexed, and females were culled to harvest the 1W timepoint. Fresh tissue was used for ATAC-Seq and RNA-Seq, whilst ovaries for ChIP-SICAP were snap-frozen and stored at −80°C until the day of the experiment. *Usp7^fl/fl^* mice were originally developed by Kon *et al.*(*78*) and were obtained from Novellasdemunt *et al.*(*103*). *Sf1:Cre^tg/+^*mice were developed by Bingham *et al.*(*79*).

### Generation of Foxl2^EGFP^ reporter strain

The *Foxl2^EGFP^* reporter strain was generated by the Genetic Modification Service at the Francis Crick Institute using CRISPR-Cas9 assisted targeting to insert a P2A and EGFP sequence after Exon 1 in frame of the *Foxl2* sequence, followed by a stop codon (Fig.S5A-E). The donor vector contained a 783bp insert of P2A and eGFP, with 1kb homology arms either side, synthesised by ThermoFisher. The guide sequence used was 5’-TCTCTGAGTGCCAACGCGCG-3’ and was cloned into the BbsI site of pX459 vector (Addgene #62988). The gene targeting was performed by co-transfection of pX459 and the synthetic donor vector into B6N 6.0 embryonic stem cells using Lipofectamine 2000. Clones were screened using an insertion PCR with primers within the EGFP (Table S1), followed by a long-range PCR with primers across the entire transgene and Sanger sequencing. Copy number was evaluated using digital droplet PCR (ddPCR, Fig.S5F, and one correctly targeted homozygous ESC clone was microinjected into goGermline blastocysts (*104*), which were then transferred into pseudo pregnant females. The resulting chimeras were crossed to C57BL/6J albino (B6(Cg)-Tyr^c-2J^/J) mice and offspring were validated for the correct integration of the transgene using the above screening PCRs followed by Sanger sequencing confirmation of the resulting amplicons. The ddPCR was performed for copy number evaluation of the F1 heterozygous mice and no evidence of random integrations of the transgene was found. Long read sequencing (Oxford Nanopore Technologies) was performed on two 3kb amplicons spanning the targeted insertion, generated by PCR and no aberrant on-target mutations or deletions were found 1.6kb upstream and 1.8kb downstream of the insertion.

### Immunofluorescence

Ovaries and testes were fixed after dissection by immersion in 4% PFA at 4°C. Immunofluorescent stainings were performed on cryosections. The following primary antibodies were used: rabbit anti-SOX9 (Millipore, cat. number AB5535), rat anti-GFP (Nacalai Tesque 04404-84), rabbit anti-FOXL2 (a generous gift from Dr Dagmar Wilhelm), rabbit anti-DDX4 (Abcam ab27591). Slides were then incubated with Alexa Fluor secondary antibodies. Imaging of stained tissue sections was performed on a Leica SPE confocal microscope.

### High Resolution Episcopic Microscopy (HREM)

*Usp7^fl/fl;^Sf1:Cre^tg/+^* mutant and *Usp7^fl/fl;^Sf1:Cre^+/+^*control ovaries were collected in PBS and immersed in Bouin’s fixative for a minimum of 12h at 4°C. Fixed tissue was extensively washed in PBS followed by dehydration in a series of graded methanol and 48h incubation in a mix of JB-4/Eosine/Acridine orange to allow proper sample infiltration. Samples were embedded in fresh JB-4/Eosin/Acridine orange mix containing accelerator for polymerization (*105, 106*). Embedded blocks were sectioned on a commercial oHREM (Indigo Scientific) at 0.70µm and sequential images of the block surface were acquired under GFP excitation wavelength light using Olympus MVX10 microscope and high-resolution camera (Jenoptik). The Eosin gives the resin a fluorescent spectrum close to that of GFP, and it is the differential quenching of this depending on the nature of the tissue that gives a negative image of the sample. After acquisition, the stack was cropped to accommodate the sample volume and adjusted for grey level using Photoshop. Data were processed for isotropic scaling, orthogonal re-sectioning and 25% downscaling with tailor made scripts. 3D volume rendering was produced using Osirix and/or Horos.

### Tissue collection and chromatin preparation for ChIP-SICAP

Ovaries from E14.5, 1W and 8-weeks old C57BL/6 mice were dissected in PBS, snap-frozen, and stored at −80°C. A pool of 140 embryonic E14.5 gonads, 30 1W ovaries, and 5 8W ovaries were used for each replicate. Chromatin was prepared using a modified version of the Active Motif High Sensitivity Chromatin Prep Kit protocol. Frozen tissue was pulverised under liquid nitrogen using mortar and pestle Bel-Art™ SP Scienceware™ Liquid Nitrogen-Cooled Mini Mortar. Ovaries were pooled to reach a minimum weight of 10 mg allowing a chromatin yield of at least 10 µg per replicate. Samples were fixed in 1 ml of Fixation Buffer (1.5 % methanol-free formaldehyde in 1% PBS) for 15 minutes on a roller at room temperature. Fixation was stopped using Stop buffer by Active Motif and incubating for 5 minutes. Chromatin preparation continued as per Active Motif protocol except for the sonication steps which were performed in a Bioruptor® Plus sonication device by Diagenode with the following settings: 40 cycles of 30”ON/30”OFF, power “high”, constant temperature of 4°C.

### ChIP-SICAP and proteomics analysis

ChIP-SICAP experiments were performed in parallel using 2 biological replicates for each timepoint. ChIP-SICAP (*18*) allows the identification of chromatin-bound proteins that colocalise with a bait protein (FOXL2 in this study) on DNA. A total of 10 µg of sonicated chromatin was used for each immunoprecipitation using 3µl of FOXL2 antibody kindly donated by Dr Dagmar Wilhelm. The original ChIP-SICAP protocol (*18, 19*) was optimised to work on frozen tissue samples as follows. Chromatin was prepared with Active Motif chromatin preparation kit as described above. After overnight immunoprecipitation at 4°C with a FOXL2-specific antibody, immune-protein complexes were captured on protein G Dynabeads. DNA was biotinylated by TdT in the presence of biotin-11-ddUTP and biotin-11-ddCTP as well as by Klenow3’exo in the presence of Biotin-7-dATP and eluted. Protein-DNA complexes were captured with protease-resistant streptavidin beads (*107*), and proteins were digested overnight at 37°C with 300 ng of LysC. Streptavidin beads carrying DNA were separated using a magnet to be processed for ChIP-seq library preparation. The supernatant, containing the digested peptides was collected and digested further by 8 hours of incubation with 200 ng trypsin. Digested peptides were cleaned using the stage-tipping technique. Briefly, digested samples were acidified by the addition of 2 µl of TFA 10%. For each sample, 50 µL of 80% acetonitrile/0.1% formic acid was aliquoted in a LCMS Certified Clear Glass 12 x 32mm Screw Neck Total Recovery Vial (Waters). ZipTip with 0.6 µL C18 resin was pre-treated by pipetting 100% acetonitrile twice by aspirating, discarding, and repeating. The ZipTip was equilibrated by pipetting 0.1% TFA, three times by aspirating, discarding, and repeating. Acidified samples were then pipetted 10 times with ZipTip to allow peptides to bind to the polymer within the tip. Following tip wash in 0.1% TFA, peptides were eluted in 80% acetonitrile/0.1% formic acid in the glass vial and the eluent dried in a speed vac. Peptides were reconstituted in 8 µL of 2% DMSO/0.1% formic acid. Peptides were separated on a 50 cm, 75 µm I.D. Pepmap column over a 70min gradient to be injected into the mass spectrometer (Orbitrap Fusion Lumos) according to the universal Thermo Scientific HCD-IT method. The instrument ran in data-dependent acquisition mode with the most abundant peptides selected for MS/MS by HCD fragmentation. The raw data were analysed using MaxQuant 2.0.1.0. The spectra were searched using the Swissprot *Mus musculus* database. Variable modification included Methionine oxidation and N-terminal acetylation. Fixed modification included Cystenine carbamidomethylation. Trypsin and LysC were chosen as the enzymes, and maximum 2 missed cleavages were allowed. Peptide and protein false identification rate was set at FDR < 0.01. The quantification values were exported to R studio to analyse significantly enriched proteins using t-test by limma package to determine Bayesian moderated t-test p-values and Benjamini-Hochberg (BH) adjusted p-values and log2FC. We considered proteins with mean fold enrichment >2 (log2FC > 1) and adj. p value <0.1 as enriched proteins.

### DNA purification for ChIP-SICAP-seq samples

Protease-resistant streptavidin beads carrying DNA samples from the above ChIP-SICAP were processed for DNA purification. SDS buffer and proteinase K (20mg/ml) were added to resuspend the beads followed by incubation for 30 minutes at 55°C. The beads were then heated at 80°C for two hours to reverse the cross-linking. A magnetic rack was used to separate the beads from the DNA, now in the supernatant. Supernatant was collected and 50 µL of Ampure beads used to purify the DNA. Purified input (no SICAP) and ChIP-SICAP DNA samples were used to generate libraries. NEB Ultra II DNA kit was used. Samples were sequenced on the NovaSeq 6000 system. Paired-end sequencing was performed (30 million reads).

### Ovary dissociation for fluorescence-activated cell sorting (FACS)

Ovaries were rinse in 1x PBS, then dissociated using add 450 µl 0.05% trypsin and 50 µl of 2.5% collagenase. Tissue was incubated at 37°C for 6 minutes (for E14.5, up 20 minutes for 1W and 8 W ovaries after mincing). Enzymatic reaction was quenched by adding 200 µl PBS/3% FBS. Supernatant was removed without disrupting the gonads, then tissue clumps were resuspended in 300 µl PBS/3% FBS and gently pipetted up and down until gonads were disaggregated. Cell mixture was passed through a 30 µm filter cap into a sorting tube. An extra 200 µl of PBS/3% FBS was used to rinse the tube and filtered. Samples were sorted with a FACS Aria III to exclude debris (side scatter versus forward scatter) and doublets (area versus width) and to select only live cells by excluding cells positive for DAPI. Cells were collected directly into a low-retention 1.5 ml tube with 200 µl PBS/3% FBS and kept on ice.

### RNA extraction, library preparation and sequencing

A total of 50000 GFP positive cells were sorted, and RNA was extracted using the RNeasy Plus Micro Kit (cat. no. 74034) by Qiagen. RNA quality was assessed by Bioanalyzer and samples with a RIN value>8 were processed for library preparation. A total of 10ng of RNA was used to prepare libraries using the NEBNext® Low Input RNA Library Prep Kit (E6420L). Samples were subsequently sequenced on the NovaSeq 6000 system, at a dept of 25 million reads (SR75) per sample.

### ATAC-Seq, library preparation and sequencing

GFP positive cells isolated by FACS from gonads at E14.5, 1W and 8W (three biological replicates per timepoint) were used for ATAC-Seq library preparation using Active Motif ATAC-Seq Kit (cat. number 53150). In brief, 100000 GFP positive sorted cells were washed in PBS, pelleted, and lysed in ATAC Lysis Buffer to isolate intact nuclei. Tagmentation of nuclei was performed for 30 minutes in a thermomixer at 37°C set at 800 rpm. Tagmented DNA was purified using DNA purification columns and library amplification was performed using Illumina’s Nextera adapters. A unique combination of i7/i5 primers was used to allow for multiplexing and sequencing on the same flow cell. PCR amplification included 5 minutes incubation at 72°C, followed by 30 sec of DNA denaturation at 98°C and 10 cycles of: 98°C for 10 sec, 63°C for 30 sec and 72°C for 1 minute. PCR products were cleaned up using SPRI beads and size distribution of PCR amplified libraries was assessed on Tapestation. Samples were subsequently sequenced on the NovaSeq 6000 system.

### mRNA-Seq alignment and quantification

Reads were processed using the publicly available nf-core(*108*) rnseq pipeline v3.3 with the STAR(*109*)/RSEM(*110*) option against mouse genome assembly GRCm38 and Ensembl release 81 transcript annotations. Other options were left as default. Gene-level RSEM abundance estimates were imported into R using the Bioconductor package tximport’s tximport function (*111*) for further analysis using DESeq2(*66*). Normalisation factors were calculated per gene using the default DESeq2 function, correcting for library size and feature length. Variance-stabilised abundance estimates were calculated using the VST function. PCA analysis was performed on the variance stabilised values using all available genes.

### mRNA-Seq differential expression analysis and clustering

Differential expression between treatment groups was assessed using a DESeq2’s Wald test in a pairwise manner, with false-discovery rate control based on an independent hypothesis weighting (IHW)(*112*). Shrunken log2 fold changes were calculated using the lfcShrink function (type=”ashr”). Significance was assessed based on a combined FDR<0.01, a shrunken log2FC>2 and a baseMean (i.e. mean abundance across all samples)>2. A combined list of significantly differentially expressed genes from all pairwise comparisons was created and used to generate a heatmap showing how patterns of expression changed across samples. Per-gene z-scores were calculated from the variance stabilised abundance estimates. K-means clustering was applied (km=7) to split the data into slices of similarly changing genes. Within each slice genes were clustered using a Euclidean distance metric and complete-linkage.

### Gene Ontology enrichment analysis

Gene Ontology (GO) Biological Process (BP enrichment analysis of the individual gene lists and the kmeans clusters was calculated relative to a genomic background via the compareCluster function from the clusterProfiler Bioconductor package(*113*) (fun=“enrichGO”, OrgDb=“org.Mm.eg.db”, ont=”BP”, pAdjustMethod=“BH”, pvalueCutoff=0.05, qvalueCutoff=0.01, minGSSize=10, maxGSSize=500).

### ChIP-SICAP: ChIP-Seq read alignment, peak calling and peak annotation

Reads were processed using the publicly available nf-core (*108*) chipseq pipeline v1.2.2 against the mouse genome assembly GRCm38 (--narrow_peak, --single_end, --deseq2_vst, min_reps_consensus 2, --macs_fdr 0.05), applying the Encode GRCm38 blacklist definitions to filter out problematic regions. A peak was only considered to be genuine if present in both biological replicates for each treatment group. A consensus peak set was then constructed by combining the peak calls from the individual treatment groups into a set of non-overlapping intervals. Consensus peaks were annotated using the Bioconductor package ChIPpeakAnno’s annotatePeakInBatch function (*114*) against the protein-coding gene component of Ensembl release 81. Gene assignments were made based on peak-centre minimal proximity to a gene’s TSS(output=“nearestLocation”,multiple=FALSE,maxgap=1L,PeakLocForDistance=“middle”,FeatureLocForDistance=“TSS”,select=“first”,bindingType =“fullRange”). The distribution of peaks mapping to genomic features was similarly assessed using ChIPpeakAnno’s “genomicElementDistribution” function, defining promoter regions as +/-2kb around a TSS and downstream regions as the 2kb region downstream of genic boundaries.

### DiffBind analysis

Consensus peak occupancy analysis was conducted using the Bioconductor package DiffBind (*115*). Briefly, reads were counted from the nfcore chipseq pipeline alignment files (BAM) over a combined set of consensus peaks (intervals). Control (input) read counts were subtracted from the targeted samples at each interval: dba.count (samples.dba, peaks=consensus.gr, summits=FALSE,filter=0,bSubControl=TRUE,minCount=0,bUseSummarizeOverlaps=TRUE,m apQCth=0). The resulting DiffBind object was normalised using a background approach, which calculates scaled factors from read counts against a set of 15000bp genomic bins considered large enough not to show differential enrichment across samples: dba.normalize (samples.dba,background = TRUE, method=DBA_DESEQ2). Normalised peak counts for all peaks were subsequently used for Pearson correlation analysis and PCA analysis. Differential binding affinity analysis was conducted in a pairwise fashion between treatment conditions using DESeq2(*66*). A FDR of <0.05 was used to threshold significant changes in peak occupancy. Peak showing significant changes in occupancy were submitted for read-depth profiling in the region +/-1.5kb from their centres. Plots are presented at the merged replicate level, stratified based on gain/loss status (i.e. direction of fold change).

### Clustering of ChIP-Seq peaks

The normalised peak occupancy estimates were used to cluster the consensus peaks into sets that showed similar patterns of variation in occupancy across conditions. Peak estimates were merged across replicates using a mean, a pseudo-count of 0.1 was added prior to a log2 transformation. Per-peak z-scores were calculated. K-means clustering was applied (km=7, 1000 repeats) to split the data into slices of similarly behaving peaks. Within each slice peaks were clustered using a Euclidean distance metric and complete-linkage.

### Motif enrichment analysis

Analysis was performed using the “calcBinnedMotifEnrR” from the monaLisa package(*69*), by scanning the sequence of consensus peaks +/-500bp from their centres against the PWMs from JASPAR 2020(*116*). Significance of enrichment was assessed against a background of equivalently sized sequences randomly sampled from the genome. Sequences were weighted to correct for GC and k-mer composition differences between fore-and background sets. Statistically significant enriched motifs within each cluster were determined using a one-tailed Fisher’s exact test with Benjamini-Hochberg adjusted p-values pval<0.0001 and subsequently prioritised based on their log2 enrichment scores (descending).

## ATAC-Seq

### Read alignment and peak calling

Reads were processed using the publicly available nf-core(*108*) atacseq pipeline v1.2.1 against the mouse genome assembly GRCm38 (--deseq2_vst --min_reps_consensus), applying the Encode GRCm38 blacklist definitions to filter out problematic regions. A peak was only considered to be genuine if present in at least 2 biological replicates for each treatment group. A consensus peak set was then constructed by combining the peak calls from the individual treatment groups into a set of non-overlapping intervals.

### Peak annotation

Consensus peaks were annotated using the Bioconductor package ChIPpeakAnno’s annotatePeakInBatch function (*114*) against the protein-coding gene component of Ensembl release 81. Gene assignments were made based on peak-centre minimal proximity to a gene’s TSS: (output=“nearestLocation”, multiple=FALSE, maxgap=-1L, PeakLocForDistance=“middle”, FeatureLocForDistance=“TSS”, select=“first”, bindingType=“fullRange”). The distribution of peaks mapping to genomic features was similarly assessed using ChIPpeakAnno’s genomicElementDistribution function, defining promoter regions as +/-2kb around a TSS and downstream regions as the 2kb region downstream of genic boundaries.

### DiffBind analysis

Consensus peak occupancy analysis was conducted using the Bioconductor package DiffBind (*115*). Briefly, reads were counted from the nfcore chipseq pipeline alignment files (BAM) over a combined set of consensus peaks (intervals) dba.count (samples.dba,peaks=consensus.gr,summits=FALSE,filter=0,bSubControl=FALSE,minCount=0, bUseSummarizeOverlaps=TRUE,mapQCth=0). The resulting DiffBind object was normalised DESeq2’s RLE method(*66*) based on the peak counts dba.normalize(samples.dba,method=DBA_DESEQ2,normalize=DBA_NORM_RLE,library=DB A_LIBSIZE_PEAKREADS). Normalised peak counts for all peaks were subsequently used for Pearson correlation analysis and PCA analysis.

Differential binding affinity analysis was conducted in a pairwise fashion between treatment conditions using DESeq2(*66*). A FDR of <0.05 was used to threshold significant changes in peak occupancy. Peak showing significant changes in occupancy were submitted for read-depth profiling in the region +/-1.5kb from their centres. Plots are presented at the merged replicate level, stratified based on gain/loss status (i.e. direction of fold change).

### Data integration

Data from the RNA, ATAC and ChIP experiments were integrated at the gene level. The ATAC and ChIP data were assigned to protein-coding genes based on minimal peak proximity to a TSS. Where this resulted in multiple peaks being assigned the same gene, the one closest to the TSS was given precedence. Identification of stably expressed and dynamically regulated genes: genes dynamically regulated by FOXL22 were defined as the intersect of i) genes differentially expressed in at least one of the pairwise comparisons from the RNA-seq analysis and ii) bound by Foxl2 based on the presence of a ChIP-seq peak in at least 1 timepoint. Stably expressed genes were defined as those bound by FOXL2 but now showing differential expression. Genes not bound by FOXL2 but showing differential expression were not considered to be directly regulated by FOXL2. Stably expressed genes were subsequently divided into “activated” and “repressed” subsets based on the variance stabilised (VST) normalised counts from the RNA-seq analysis. A VST score of >=8.3 in at least 1 sample was enough to quality a gene as “activated”, while all others were regarded as “repressed”.

### Cytoscape visualisation of FOXL2 protein interactors

GO enrichment analysis of FOXL2 interactors identified by ChIP-SICAP and fold changes over no-antibody control in Table S3 were loaded onto Cytoscape v.3.9.1(*117*). Proteins were clustered based on their function. The cytoscape app enhancedGraphics(*118*) was used to overlay bar charts depicting the fold changes over control of each protein in each timepoint.

**Fig. S1.**
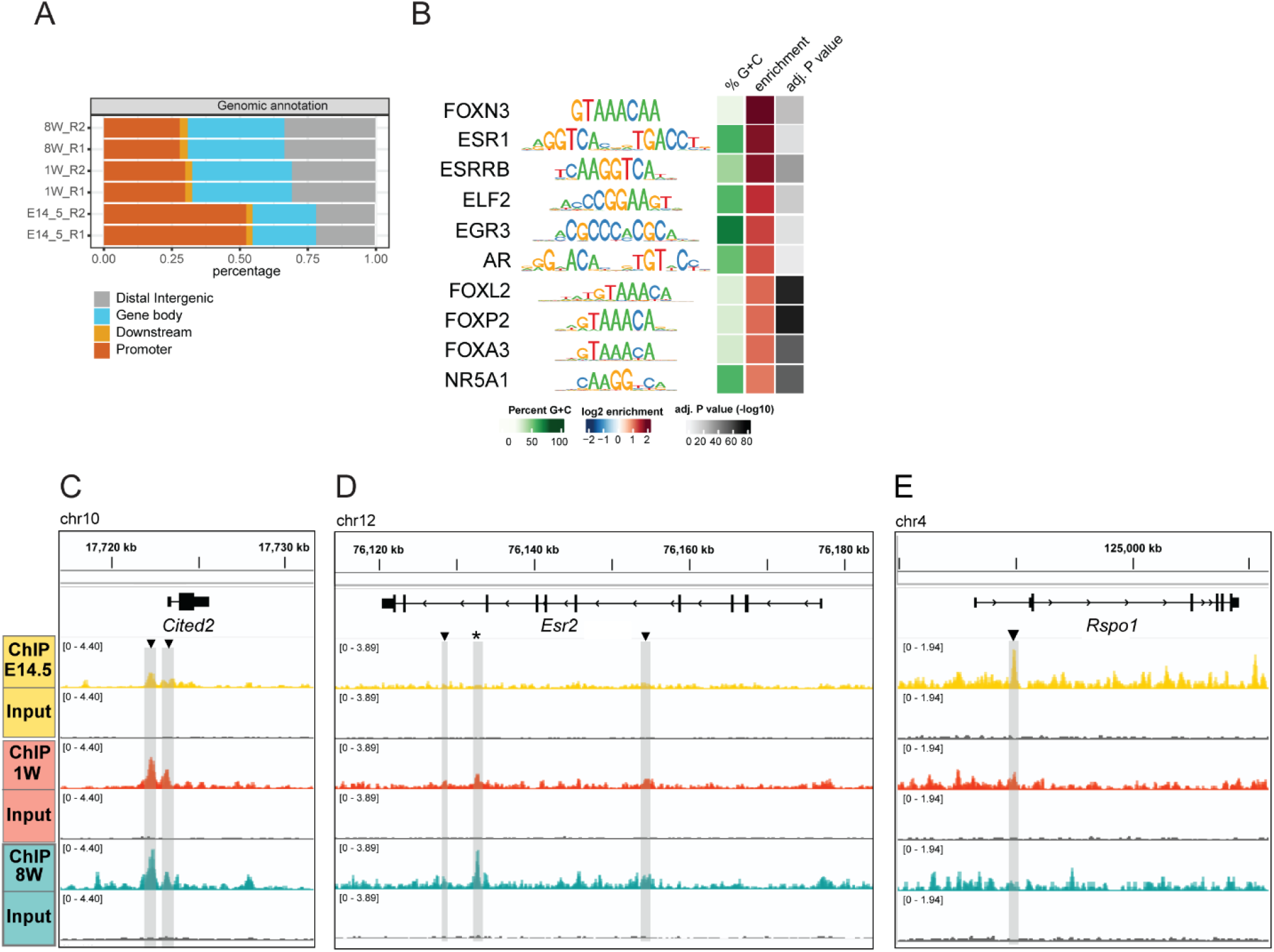
FOXL2 genome-wide occupancy across ovarian development. (A) Annotation of consensus peaks by ChIPpeakAnno (B) Enrichment analysis of transcription factor binding sites associated with consensus peaks significantly changing across the timecourse. Analysis performed using monaLisa package (*119*). Top 10 representative TF are shown (*p-value*<0.0001). (C-E) IGV tracks representative of FOXL2-bound regions. Main tracks show normalised read-depth coverage. Grey bars and black arrows highlight significant peaks over input. (C) *Cited2*, (D) *Esr2*(*39*), and (E) *Rspo1.* Star denotes known *Esr2* enhancer (*39*).

**Fig. S2.**
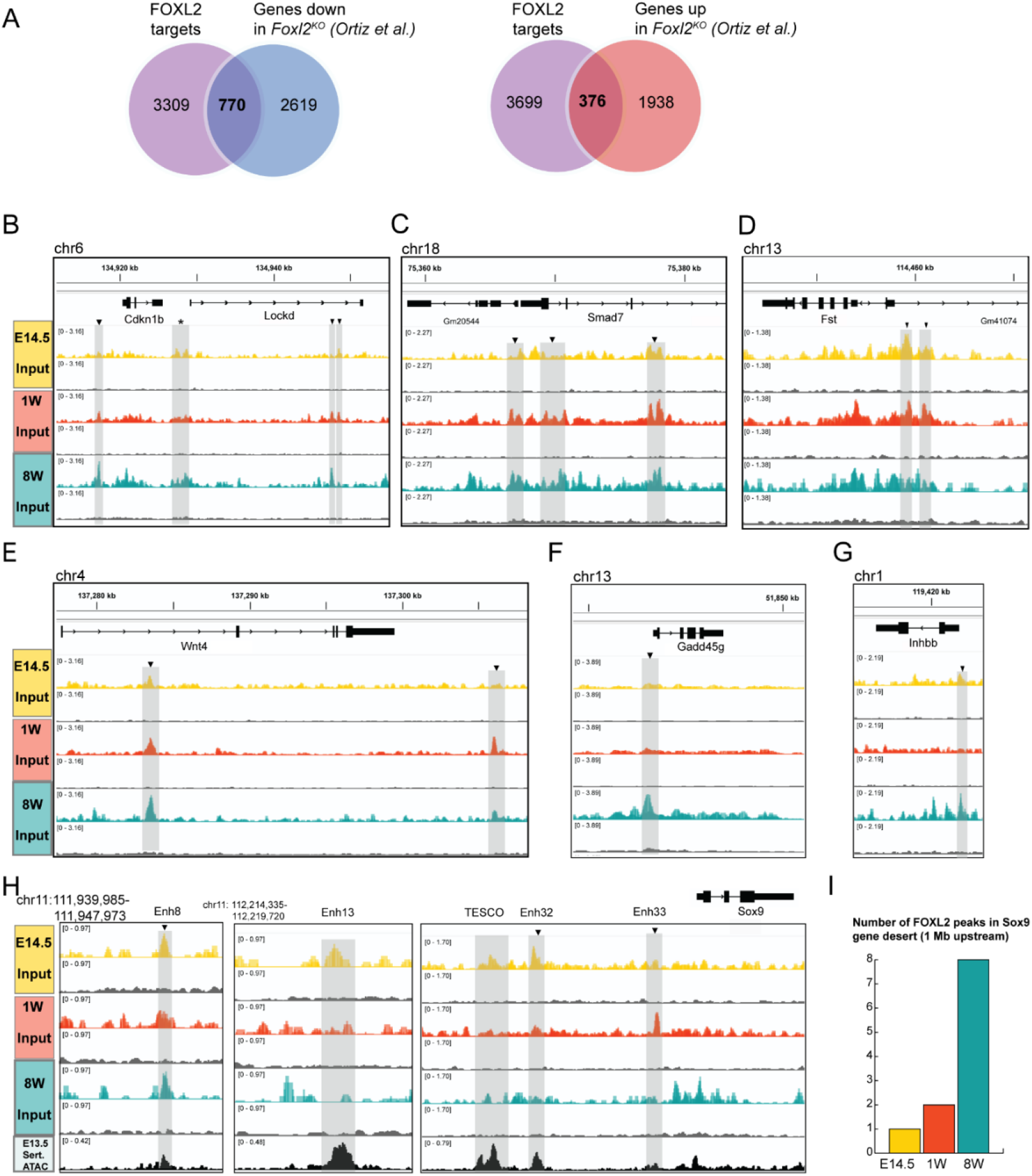
FOXL2 genome-wide occupancy across ovarian development. (A) Venn diagram showing the overlap between FOXL2 target genes identified at any point throughout ovarian development (E14.5, 1W and 8W) and total of genes downregulated/upregulated in *Foxl2^-/-^* null mutant ovaries collected at 13.5, 16.5 dpc and birth compared to wildtype controls (Ortiz *et al.*(*28*) and Data S2). Genomic overviews of FOXL2 peaks in representative genes likely to be activated by FOXL2 including *Cdkn1b* (B), *Smad7* (C), and *Fst* (D). Representative genes likely to be downregulated by FOXL2, including *Wnt4* (E), *Gadd45g* F), *Inhbb* G) and *Sox9* (H). (I) Bar chart displaying the total number of significant peaks identified in each timepoint in the 1Mb gene desert upstream of *Sox9*.

**Fig. S3.**
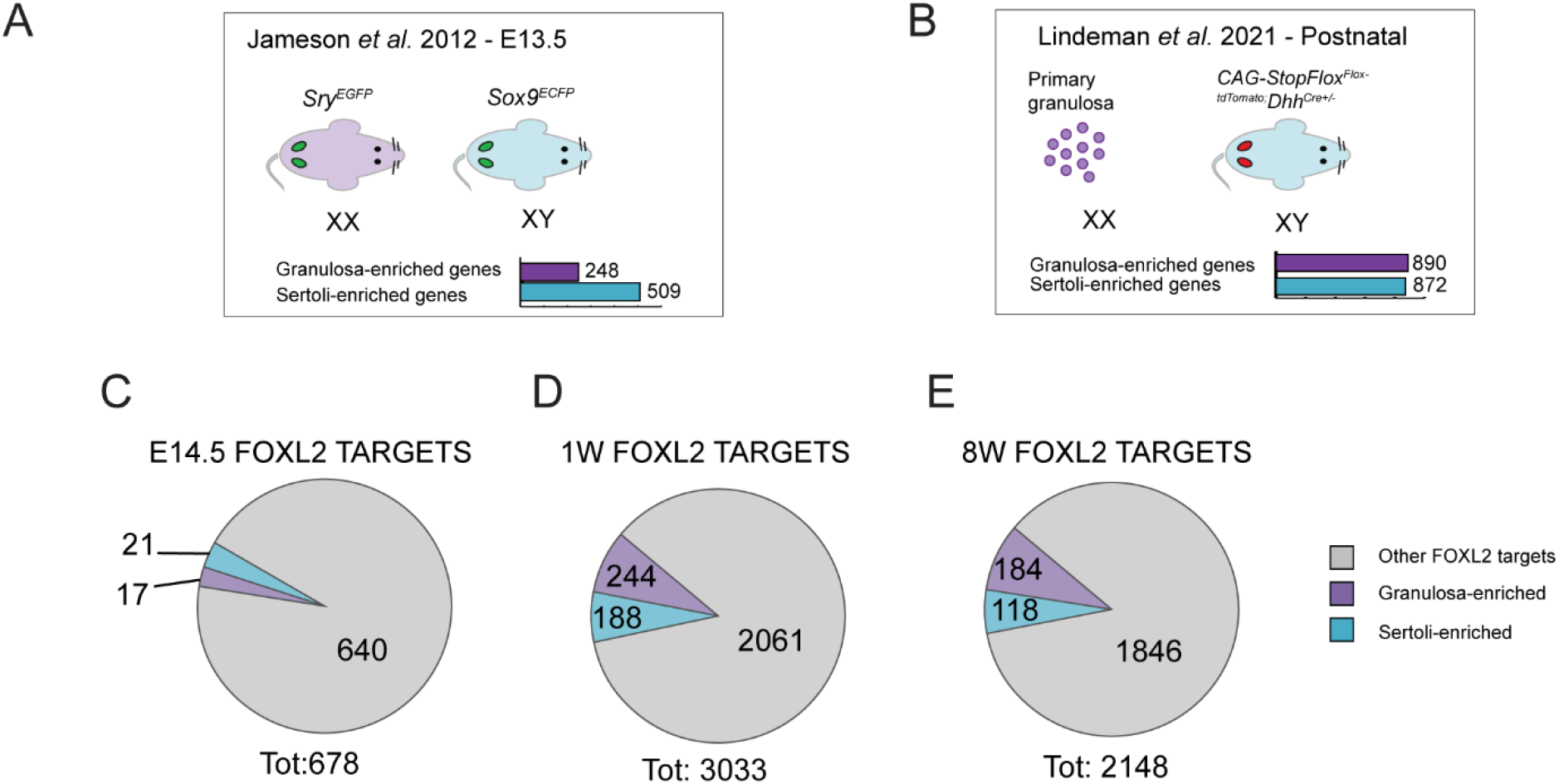
The contribution of FOXL2 to the regulation of granulosa and Sertoli-enriched genes is greater at postnatal stages. (A-B) Overview of datasets used: microarray analysis performed on XX and XY E13.5 gonads, by Jameson *et al*. (*49*), and RNA-Seq analysis by Lindeman *et al.*(*50*) of primary granulosa cells isolated from 23-29 days old mice, compared to Sertoli cells from P7 XY pups. Bar charts indicate the number of genes enriched in either granulosa (purple) or Sertoli cells (blue). (C) Pie chart depicting the proportion of FOXL2 target genes, as identified at E14.5, and classified as either granulosa-or Sertoli enriched at E13.5(*49*). (D) Proportion of FOXL2 target genes, as identified at 1W and (E) 8W, classified as either granulosa-or Sertoli-cell specific and are enriched postnatally. Grey denotes other FOXL2 target genes identified by our ChIP-SICAP and not overlapping with the lists of granulosa/Sertoli-enriched genes from the two studies.

**Fig. S4.**
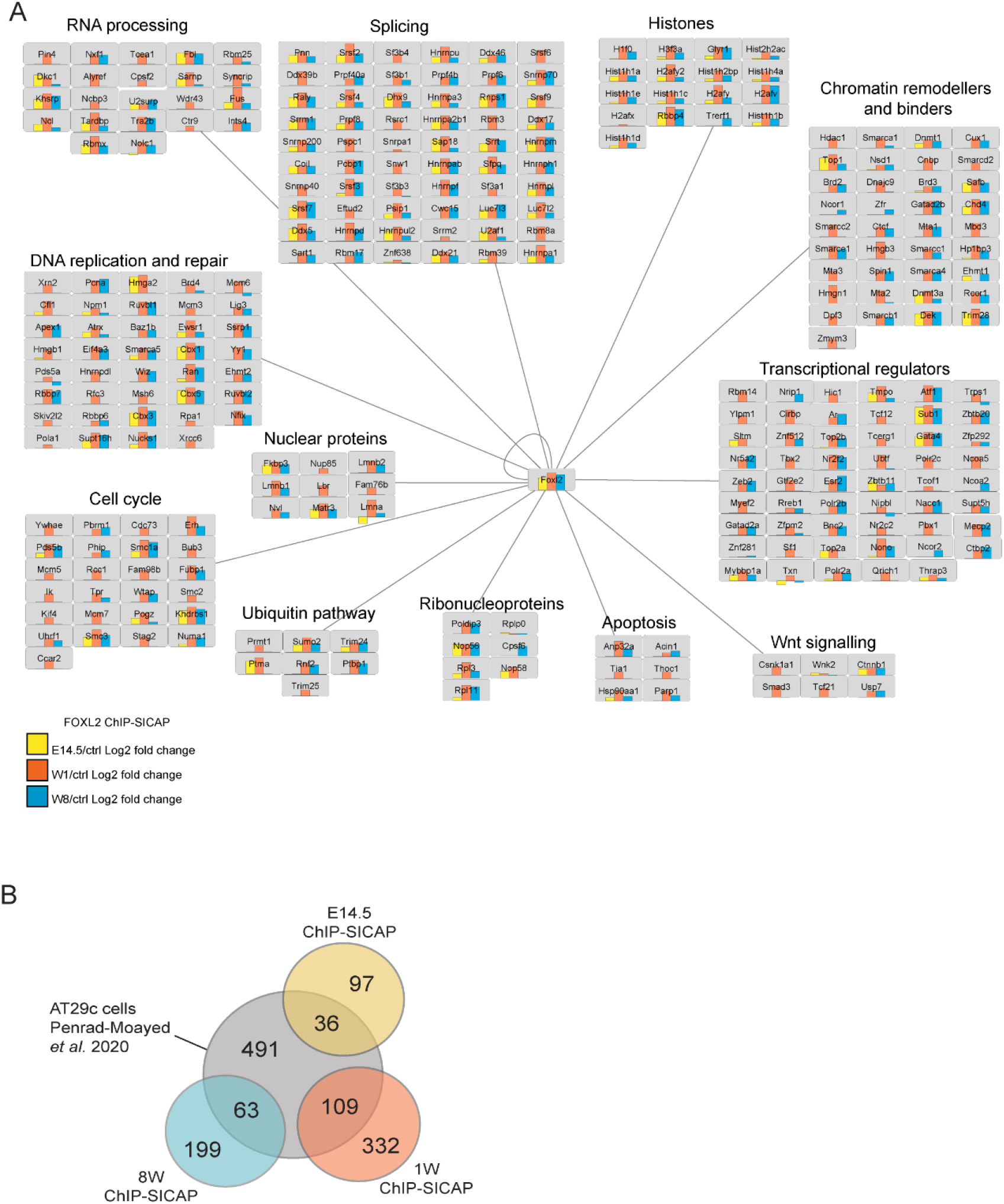
Cytoscape network of FOXL2 interactome across ovarian development. (A) Proteins interacting with FOXL2 on-chromatin were clustered by GO Processes and visualised with Cytoscape. Enrichment values over no-antibody control were depicted as barcharts (yellow=E14.5, orange=1W, blue=8W, n=2, fold change over no antibody control >2 in at least one timepoint, adj-*pvalue*<0.1) and visualised within the network using enhancedGraphics Cytoscape plugin (*118*). (B) Overlap of protein interactors found in our study compared to a cell line (AT29c) whole proteome study by Penrad-Mobayed *et al.*(*52*).

**Fig. S5.**
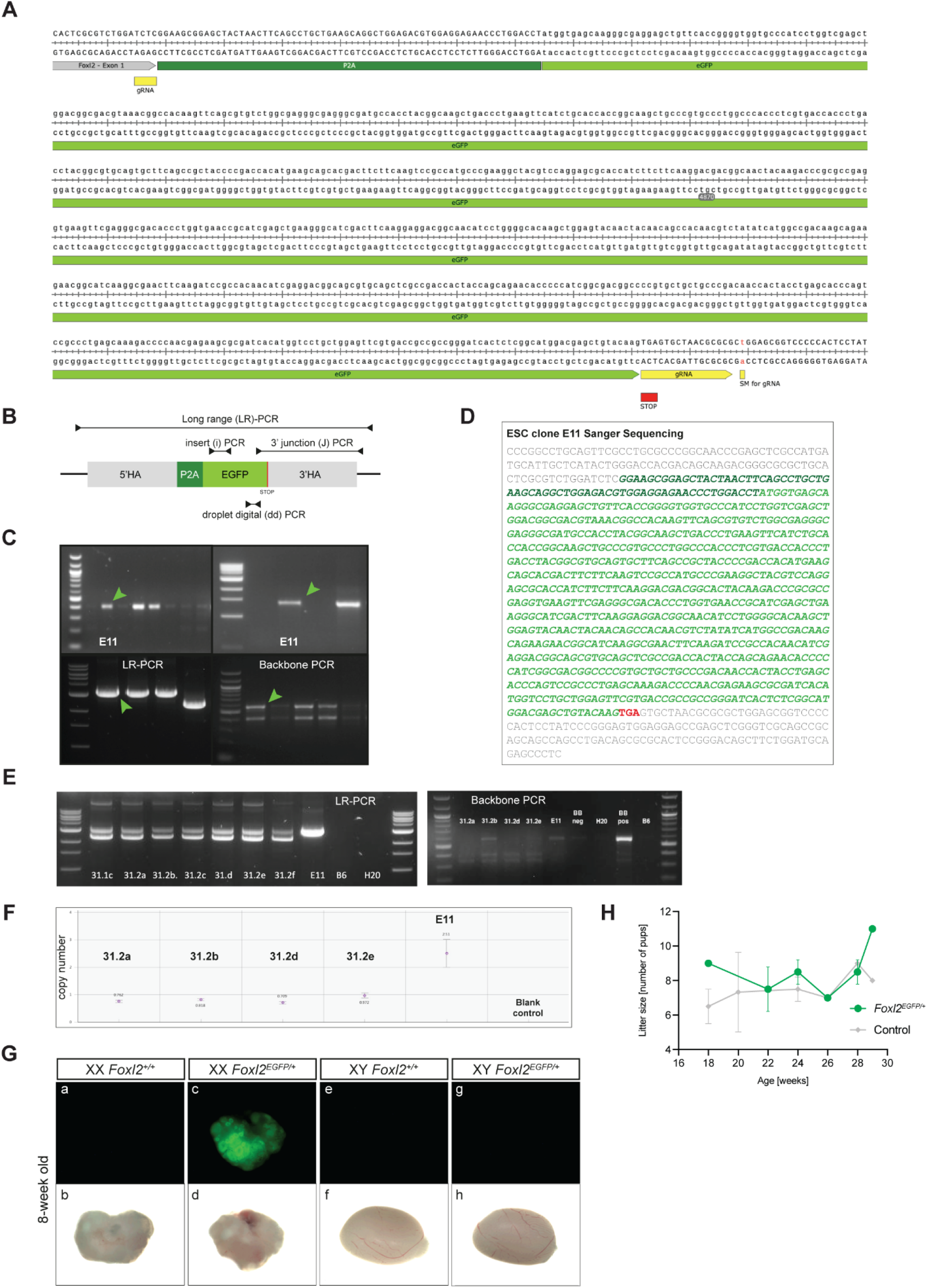
Design of *Foxl2^EGFP^* construct, genotyping strategy, and characterisation of the *Foxl2^EGFP^* mouse line. (A) Design of gRNA (yellow box) and associated silent mutation (SM) in PAM sequence as well as 783bp P2A-EGFP construct (green boxes). (B) Location of primers for insert, junction, long-range and digital droplet PCR. (C) ES cell screening identified clone E11 (green arrows), which showed successful EGFP integration depicted by a single band in insert, junction, and long-range PCR. (D) Correct integration of EGFP into Foxl2 locus of clone E11 was confirmed via sanger sequencing. (E) Long-range PCR of chimeric offspring as well as clone E11 confirmed heterozygous and homozygous EGFP insertion, respectively. No evidence of backbone integration was found. BB pos= backbone positive; BB nrg= backbone negative. (F) Copy number evaluation of chimeric offspring as well as ES cell clone E11 by ddPCR confirmed absence of random transgene integration. (G) Representative images of ovaries and testes dissected from adult mice (8 weeks) and imaged with a fluorescence microscope. Upper row: GFP channel, lower row: brightfield channel. Adult mouse ovaries wildtype (a-b) and heterozygous (c-d), as well as testes wildtype (e-f) and heterozygous (g-h) are shown. Scale bars=100 µM. (H) Fertility assessment of *Foxl2^EGFP/+^* females compared to wildtype controls (C57BL/6J stock mice).

**Fig. S6.**
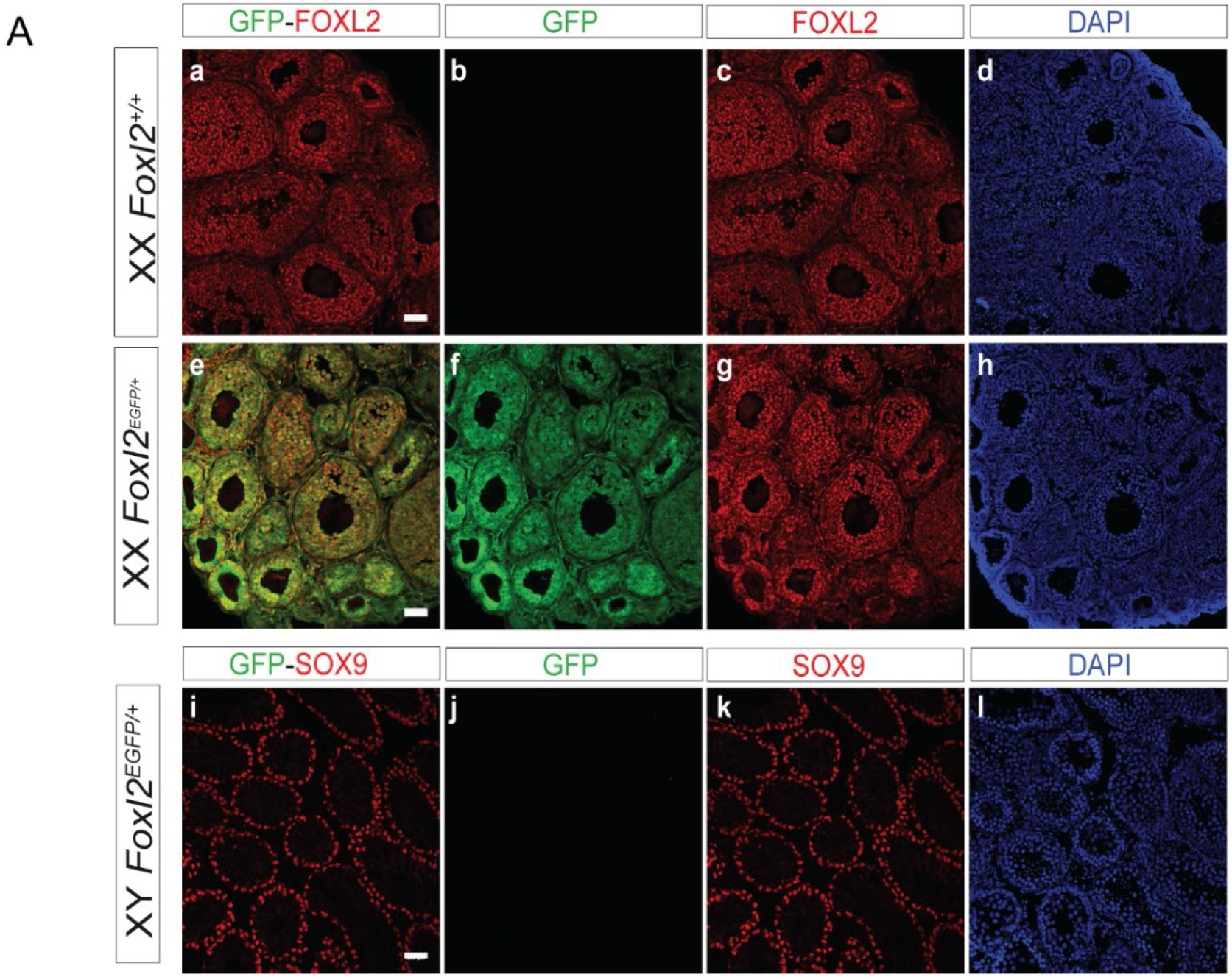
Characterisation of the *Foxl2^EGFP^* mouse line. **(A)** Immunofluorescence staining of EGFP (green) and endogenous FOXL2 (red) in adult XX *Foxl2^+/+^* ovaries (a-d), XX *Foxl2 ^EGFP/+^* ovaries (e-h), and of EGFP and endogenous SOX9 (red) in XY *Foxl2 ^EGFP/+^* testes (i-l). Scale bars: 100 µm.

**Fig. S7.**
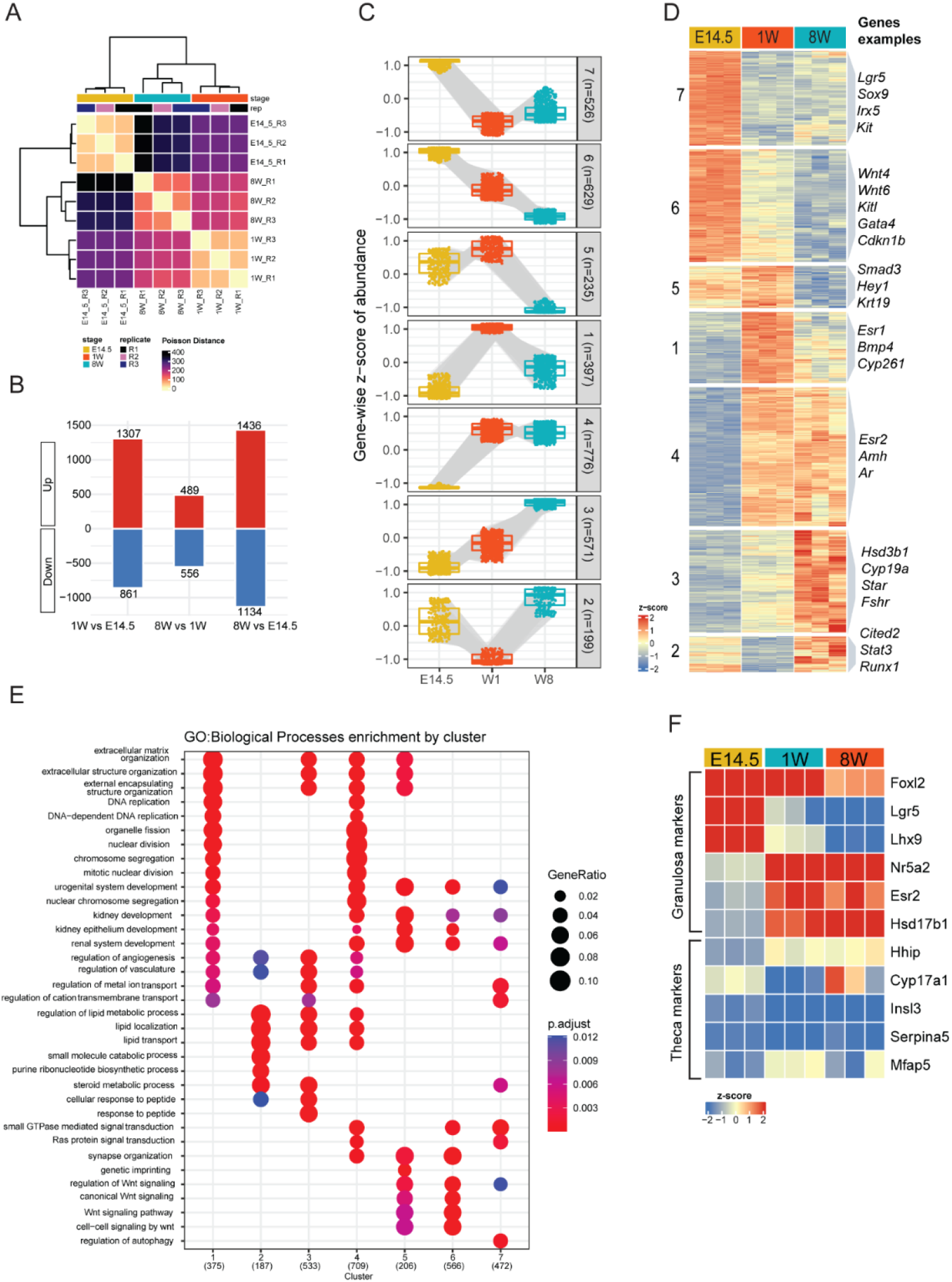
Overview of RNA-Seq analysis of *Foxl2^EGFP/+^* cells isolated throughout ovarian development. (A) Poisson dissimilarity metric to assess RNA-Seq sample similarity, n=3 biological replicates. (B) Bar plot summary illustrating the number of differentially expressed genes (DEG) identified in each pairwise comparison (*pvalue*<0.01, fold-change >2). (C) Boxplot illustrating the dynamics of gene expression changes granulosa *Foxl2^EGFP/+^* cells. (D) Hierarchical k-means clustering of the gene expression dynamics and representative genes on the right-hand side. (E) Top terms from the GO enrichment test (Cluster Profiler) showing the processes associated with DEGs from each of the seven clusters. (F) Heatmap of gene expression changes of granulosa/theca cell markers.

**Fig. S8.**
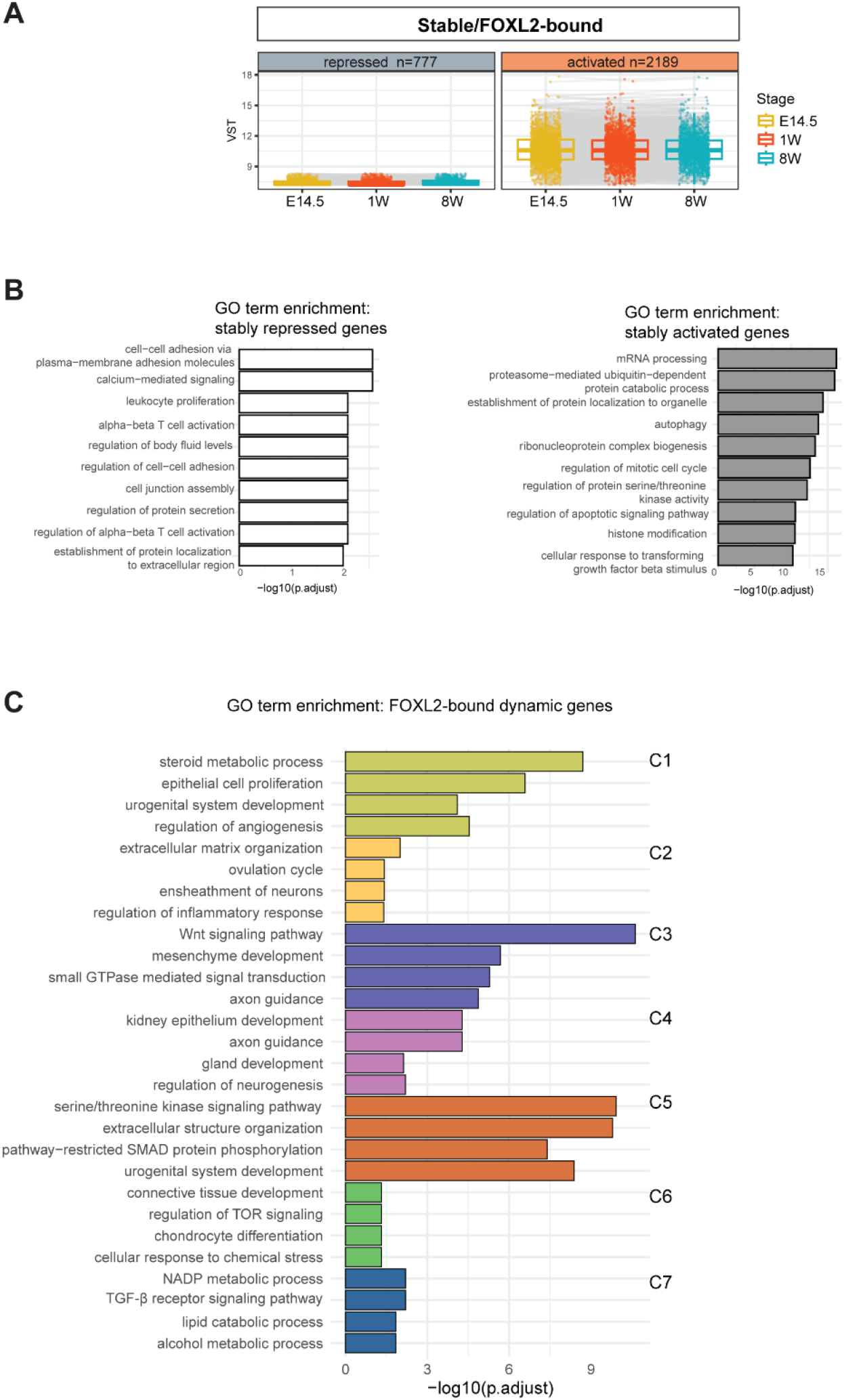
Integrative approach combining RNA-Seq and ChIP-Seq to refine the gene regulatory networks controlled by FOXL2. (A) Plot of variance stabilised (VST) gene expression values as detected by RNA-Seq on *Foxl2 ^EGFP/+^* sorted ovarian cells (n=3). (B) GO Biological processes enrichment analysis of stably repressed (left), and stably activated (right) genes. (C) GO enrichment analysis of the 1100 genes bound by FOXL2 and differentially expressed across the timecourse. Top four representative pathways are depicted.

**Fig. S9.**
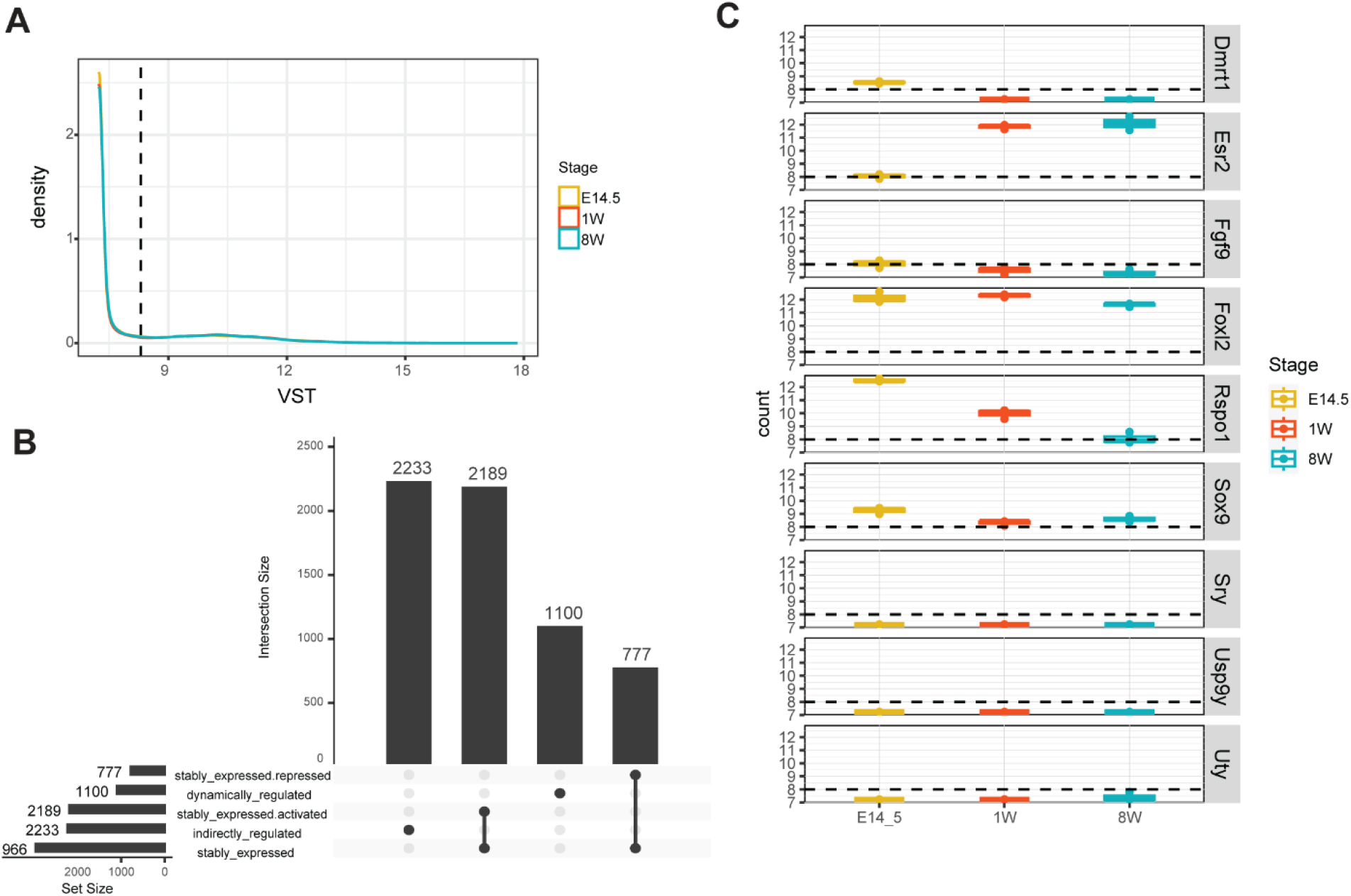
Integration of FOXL2 ChIP-Seq data with RNA-Seq of *Foxl2^EGFP/+^* cells. (A) Plot of gene density and VST cut-off =8.3 used to distinguish genes stably repressed from those stably expressed. (B) Bar plot depicting the number of genes either stably repressed or activated and bound by FOXL2. (C) Example of gene expression changes used to choose the VST cut-off. The male marker *Fgf9* was used as reference for the cut-off as known to not be expressed in the ovary.

**Fig.S10.**
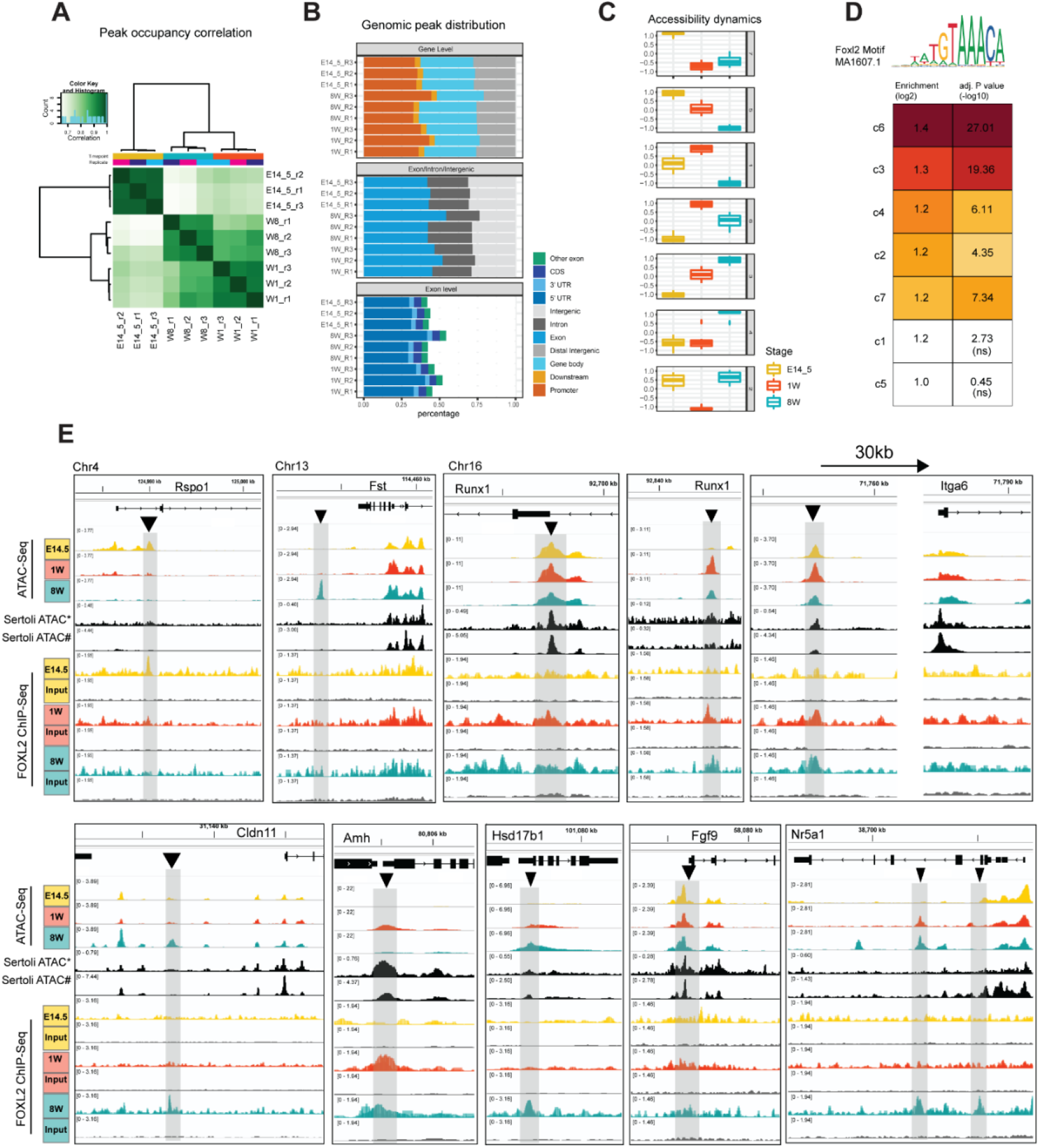
ATAC-Seq analysis of chromatin accessibility of *Foxl2^EGFP^* positive cells collected across ovarian development. (A) Poisson metric of peak occupancy correlation between timepoints. (B) Genomic annotation of consensus peaks. (C) Boxplot of z-scores representing overall trends of chromatin opening dynamics within each cluster derived from the hierarchical clustering of normalised abundancies of peaks. (D) Heatmap of enrichment scores for the FOXL2 canonical motif, ranked on enrichment score and filtered by -log10pval. Cut-off: -log10pval > 4 (i.e. pval<0.0001). (E) IGV snapshots representative of putative enhancers (grey boxes) identified within granulosa Sertoli cell marker genes, as assessed by ATAC-Seq and FOXL2 ChIP-SICAP. Black tracks show E13.5 Sertoli ATAC-Seq data re-analysed from Garcia-Moreno *et al*. (*102*), and P7 Sertoli ATAC-Seq from Lindeman *et al.* (*50*).

**Fig. S11.**
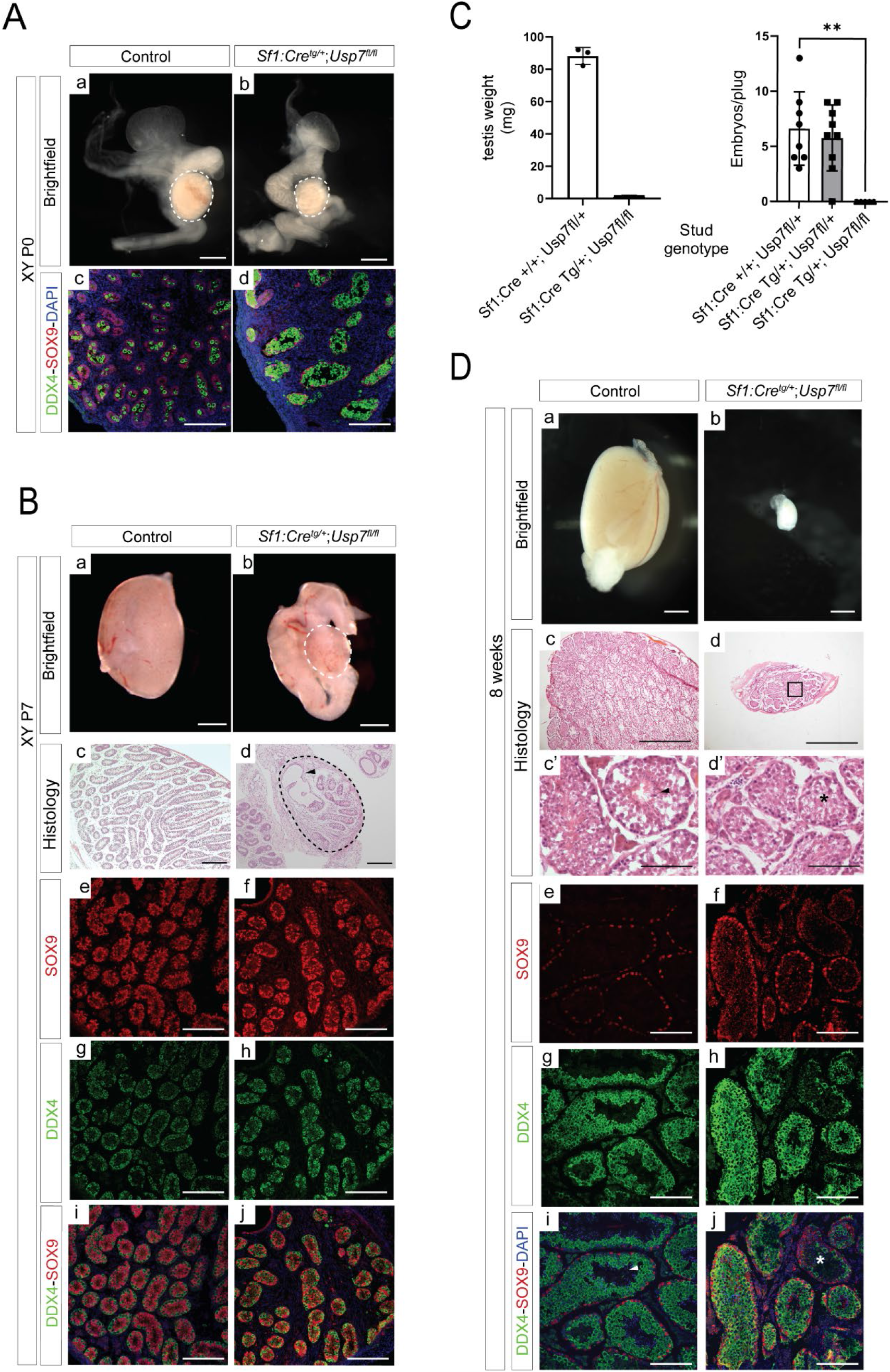
Loss of *Usp7* in Sertoli cells leads to hypogonadisms and infertility in XY mice. Characterisation of the Usp7 conditional knock-out phenotype in mouse XY testes. (A) Brightfield images showing gross morphology of control (*Usp7^fl/fl^;Sf1:Cre^+/+^*, a) and mutant testes (*Usp7^fl/fl^;Sf1:Cre^tg/+^*, b) collected at birth (P0). Scale bars = 0.5 mm in all unless otherwise specified. Images representative of n=3. Immunofluorescence analysis of germ cell marker DDX4 and Sertoli cell marker SOX9 (c-d). (B) Brightfield images showing gross morphology of control (a) and mutant (b) testes collected 1 week postnatally (P7). (c-d) Hematoxylin and Eosin (H&E) staining of section from control (c) and mutant (d) collected at P7 shows gonadal dysgenesis in mutant testes. Scale bar=200µm. (e-j) Immunofluorescence analysis of DDX4 and SOX9 at P7. (C) Left: average testis weights of adult mice in mg. Data shown are mean and standard deviation from n=3 adult mice of each genotype. Right: number of born embryos per plug for male studs of the genotyped indicated and mated with control females. Data are mean and standard deviation from n=3 *Usp7^fl/fl^;Sf1:Cre^+/+^*studs (8 plugs in total, control), from n=3 *Usp7^fl/+^;Sf1:Cre^tg/+^*studs (9 plugs in total, heterozygous mutant), and from n=3 *Usp7^fl/fl^;Sf1:Cre^tg/+^*studs (5 plugs in total, homozygous mutant). Asterisks denote p value<0.05, one-way Anova test. (D) Brightfield images showing gross morphology of control (a) and mutant (b) 8 weeks old testes. Scale bar= 1mm (a-d). (c-d) H&E staining of sections from control (c-c’) and mutant (d-d’) testes. Arrowhead in c’ points to elongated spermatids in the lumen of the seminiferous tubule. Asterisk in d’ denotes absence of these in mutants. Scale bar=100µm (c’-d’). Immunofluorescence analysis of SOX9 in control (e) and mutant (f) testis sections indicates abnormal development and organisation of Sertoli cells (SOX9, red) and impaired spermatogenesis (DDX4, germ cells, g-h). (i-j) merge. Scale bar=500µm. Images are representative of 3 biological replicates.

**Table S1.**
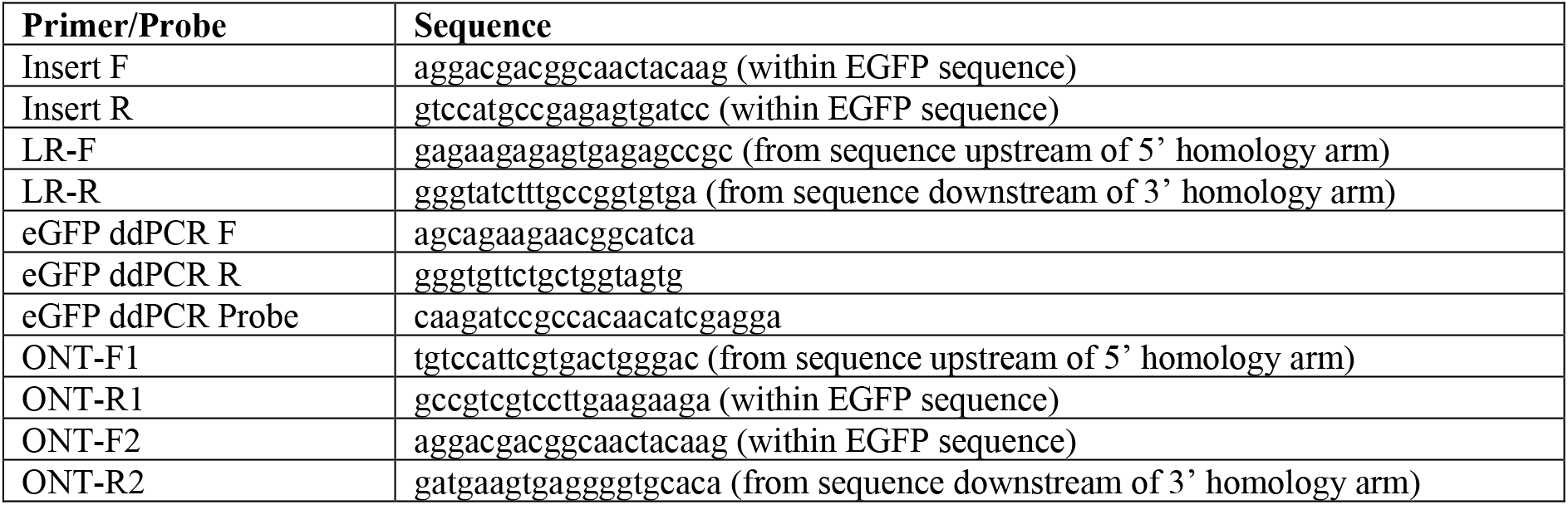
Table of primers used to genotype the *Foxl2^EGFP^* reporter strain.

### Other supplementary materials

**Movie S1.** HREM 3D rendition of adult reproductive tract from *Usp7^fl/fl^;Sf1:Cre^tg/+^* XX mice. **Data S1:** FOXL2 ChIP-SICAP analysis of genomic targets, GO of clusters and diffbind analysis. **Data S2:** Comparison of FOXL2 ChIP-SICAP with other datasets from Jameson *et al.*(*49*),

Lindeman *et al.*(*50*), and Ortiz *et al.*(*28*).

**Data S3:** FOXL2 ChIP-SICAP, proteomics dataset of FOXL2 interactors.

**Data S4:** RNA-Seq of *Foxl2^EGFP/+^* cells isolated throughout ovarian development.

**Data S5:** Data integration of ChIP-Seq, ATAC and RNA-Seq and gene-level dynamics.

**Data S6:** ATAC-Seq normalised peak counts, diffbind analysis, clustering analysis.

